# Medium-sized protein language models perform well at transfer learning on realistic datasets

**DOI:** 10.1101/2024.11.22.624936

**Authors:** Luiz C. Vieira, Morgan L. Handojo, Claus O. Wilke

## Abstract

Protein language models (pLMs) can offer deep insights into evolutionary and structural properties of proteins. While larger models, such as the 15 billion parameter model ESM-2, promise to capture more complex patterns in sequence space, they also present practical challenges due to their high dimensionality and high computational cost. We systematically evaluated the performance of various ESM-style models across multiple biological datasets to assess the impact of model size on transfer learning via feature extraction. Surprisingly, we found that larger models do not necessarily outperform smaller ones, in particular when data is limited. Medium-sized models, such as ESM-2 650M and ESM C 600M, demonstrated consistently good performance, falling only slightly behind their larger counterparts—ESM-2 15B and ESM C 6B—despite being many times smaller. Additionally, we compared various methods of compressing embeddings prior to transfer learning, and we found that mean embeddings consistently outperformed other compression methods. In summary, ESM C 600M with mean embeddings offers an optimal balance between performance and efficiency, making it a practical and scalable choice for transfer learning in realistic biological applications.

## Introduction

Machine learning (ML) has significantly advanced the field of protein biochemistry^1–4^. By leveraging large datasets and sophisticated models, ML techniques have improved the accuracy of protein property predictions^5–8^. BERT (Bidirectional Encoder Representations from Transformers) models, such as Evolutionary Scale Modeling 2 (ESM-2)^9^, ProtTrans family (ProtT5, ProtBERT, ProtAlbert)^10^, and ProtFlash^11^, are pre-trained masked protein language models (pLMs) that have proven effective at producing biologically meaningful representations of proteins. Their ability to capture evolutionary relationships from sequence data makes them particularly valuable for a wide range of downstream biological tasks. On the other hand, generative pre-trained transformer (GPT) models, such as ProGen^12^, are designed to generate novel protein sequences and have shown potential to create functional proteins. While GPT models offer great flexibility for sequence generation, BERT models excel at learning rich contextual embeddings for downstream predictions, such as predicting biological phenotypes of mutant libraries.

The pLM models learn about protein properties during pre-training, where the models are trained to predict residues that have been masked in their input sequences^13,14^. The pre-training process causes the models to encode knowledge about protein biochemistry and protein evolution in the models’ internal representations, known as embeddings, which encapsulate the biochemical characteristics of individual amino acids as well as complex higher-order interactions that reflect both local and global structural and functional properties of proteins^10,15,16^. The capacity of pLMs to learn rich representations of proteins from vast amounts of unlabeled sequence data and subsequently apply this knowledge to secondary supervised tasks, a process known as transfer learning^17^, has enabled their application in various downstream tasks including functional annotation, mutational effect analysis, and the design of novel proteins and peptides^8,18–23^. Finally, pLM embeddings can also be used for homology search, in particular when sequences are short or highly diverged^24,25^.

There has been a trend towards increasing the size of pLMs, following similar advancements in natural language processing, where model scaling laws predict that model performance systematically increases with increasing model size and commensurate increase in pre-training data^26,27^. Models such as the largest ESM-2 variant, with 15 billion parameters^16^, and more recently ESM3, with a staggering 98 billion parameters^7^, have demonstrated that scaling the model size can enhance performance by capturing more complex relationships in protein sequences. Although models such ESM3 can deliver notable gains in accuracy, their high computational cost hinders their development and broad utilization. For example, fine-tuning these large models is highly computationally demanding^28^, limiting their use primarily to private industry or highly resourced laboratories. In fact, a recent study^29^ questioned whether larger models are the best option, or if alternative training methods, such as improving data quality, increasing data quantity, or extending the number of training steps, could boost smaller models’ performance. In general, the field of AI applications in biology appears to be increasingly recognizing the benefits of smaller models. This trend is exemplified by the recent launch of the protein language model ESM-Cambrian (ESM C)^30^, which has demonstrated impressive performance in protein contact prediction, even outperforming the much larger ESM-2 15B. This shift in thinking could provide a more accessible and cost-effective alternative for the broader research community, as smaller models might offer sufficient performance for many scientific applications at a lower cost.

Here, we systematically evaluate the impact of model size on transfer learning via feature extraction, considering models with parameter counts ranging from 8 million to 15 billion across various biological datasets. These datasets include 41 deep mutational scanning (DMS) datasets^31^ and 12 different metrics calculated from proteins in the PISCES dataset^32^. Our findings suggest that the effectiveness of transfer learning is influenced by the choice of embedding compression method, dataset size, protein length, and model size. Larger models perform better with larger datasets, underscoring the need for sufficient data to maximize their utility. However, when data is limited, medium-sized models perform comparably to, and in some cases outperform, larger models. In addition, while the impact of compression methods varies depending on dataset type, mean embeddings generally yield excellent results across tasks. These findings emphasize the importance of selecting model size and embedding method based on the unique characteristics of the data and task to enhance both the efficiency and accuracy of transfer learning in protein applications.

## Results

### Mean embeddings outperform all other compression methods

Despite the general success of transformer model embeddings, their high dimensionality poses challenges in practical applications, particularly in transfer learning scenarios^33,34^. Embeddings typically need to be compressed before any downstream prediction tasks are performed. The most commonly used compression strategy is simply averaging embeddings over all sites (mean pooling)^5,19,35–37^, though other methods, such as inverse Discrete Cosine Transform (iDCT), have also been proposed^24^. Because mean pooling averages over the contributions of all sites in a sequence, this strategy may not retain all critical information, in particular in deep mutational scanning (DMS) applications, where one or a few amino acid changes can significantly affect a protein’s thermodynamic stability or enzymatic activity^38^.

To assess the effect of compression methods on transfer learning performance via feature extraction, we systematically explored a wide range of compression methods and datasets to establish a realistic and representative benchmark for evaluating pLM performance on biologically relevant tasks. First, we analyzed 40 deep mutational scanning (DMS) datasets, which primarily involve single or few point mutations relative to a reference sequence. These datasets span a wide range of experimental measurements, including genome sequencing, fluorescence, minimum inhibitory concentration (MIC), enzyme activity, protein stability, viral replication, binding assays, and microbial growth (in yeast and bacteria). They also cover proteins from diverse organisms such as humans, bacteria, and yeast. Second, we used a set of diverse protein sequences from the PISCES database, for which we computed various target variables, including physicochemical properties, instability index, amino acid frequencies, and secondary structure content. The DMS datasets test the models’ ability to predict mutation effects, while the PISCES dataset evaluates global sequence understanding.

In our initial analysis, we evaluated how embedding compression affects predictive performance in transfer learning when protein language models are used for feature extraction. In our prediction pipeline, for a given protein sequence, we extracted embeddings from the last hidden layer of the ESM-2 150M model, compressed them via one of the methods, and then used the compressed embeddings as input features in regularized regression models (LassoCV) to predict the target of interest (Figure 1). For both the DMS datasets and the PISCES sequences, we generally found that mean pooling outperformed all other compression methods we considered (Supplementary Materials, Fig. S1 and Fig. S2). However, the extent to which mean pooling performed better depended on the type of data. For diverse protein sequences, mean pooling was strictly superior in all cases, and in many cases by a wide margin (Fig. S2). By contrast, for DMS data, some alternative compression methods, including max pooling, iDCT, and PCA, were slightly better than mean pooling on some datasets, even if on average mean pooling was superior (Fig. S1). These findings suggest that mean pooling does to some extent average out effects from individual sites, yet on the whole it still tends to perform better than the alternatives.

**Figure 1.**
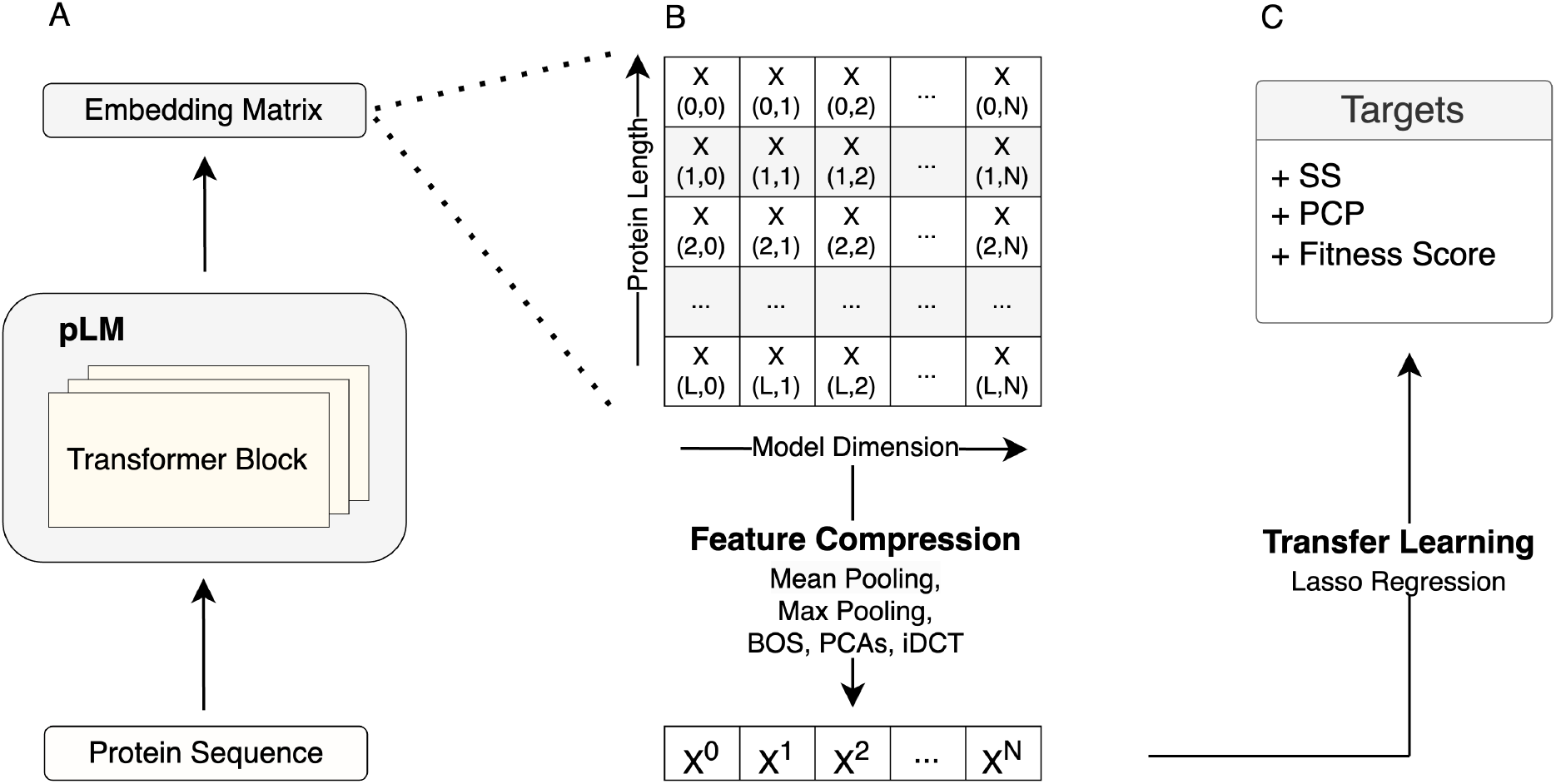
Schematic view of the transfer learning via feature extraction approach used throughout this work. A) Protein representation (embeddings matrix) extraction using the pLMs. B) Embeddings matrix compression using various methods including mean pooling, max pooling, BOS, PCAs, and iDCT. C) Transfer learning to predict downstream tasks, such as secondary structure (SS), fitness, and physico-chemical properties (PCPs).

To summarize these findings more systematically, we fit linear mixed-effects models to the results of all compression methods and either all DMS datasets or all PISCES prediction targets, respectively. In this setup, we treated the type of compression as a fixed effect and the dataset/prediction target as a random effect, to isolate the impact of the compression method while accounting for the variability between datasets. This analysis showed that mean pooling was, on average, significantly better than all other alternatives we considered, in both types of datasets (Figure 2). For DMS data, mean pooling led to an increase in variance explained (measured by *R*^2^ from the regularized regression, calculated on a hold-out test set) between 5 and 20 percentage points (Figure 2A). For diverse protein sequences, the difference was even more stark, where mean pooling led to an increase in variance explained between 20 and 80 percentage points (Figure 2B).

**Figure 2.**
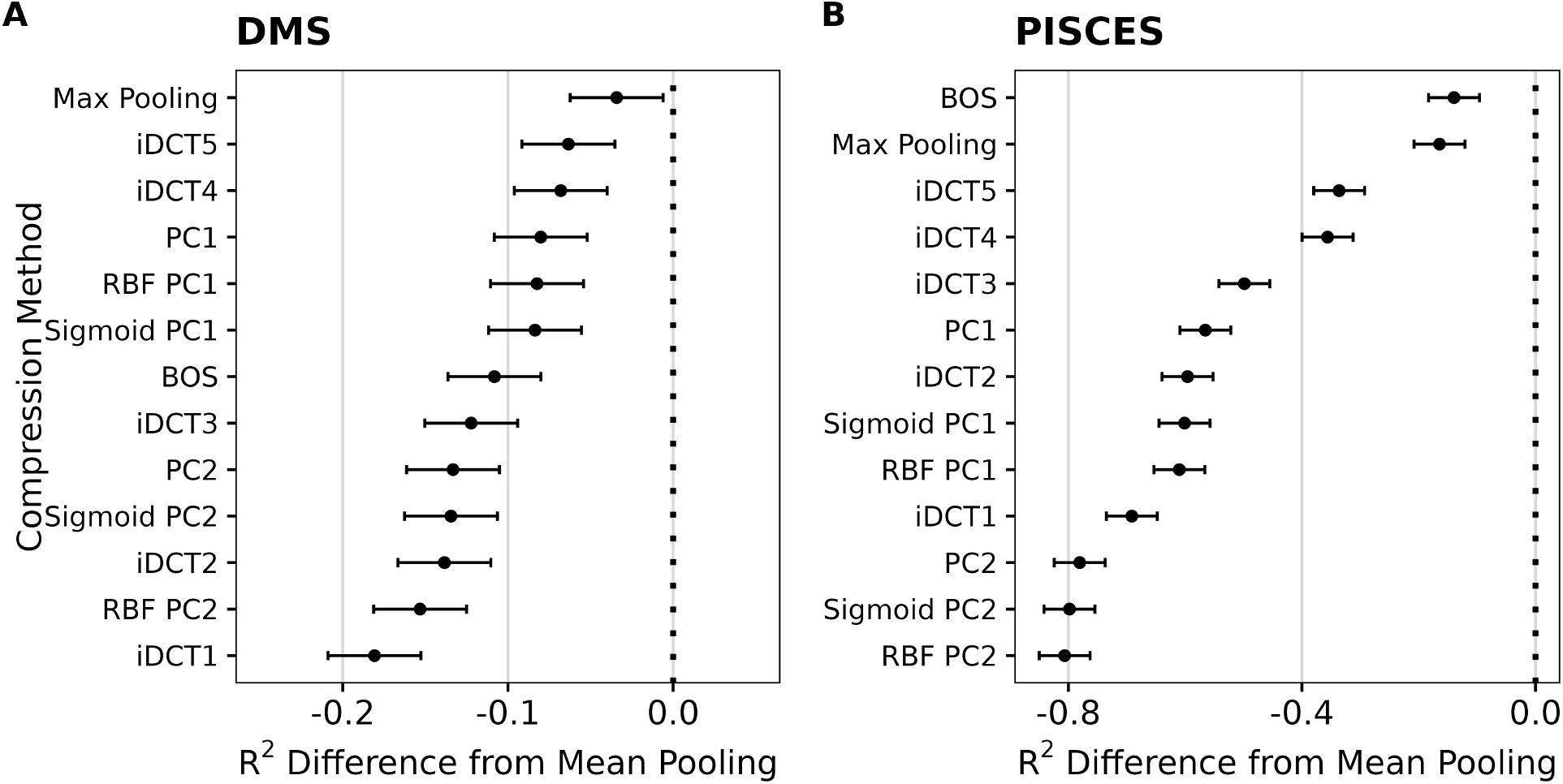
Mean reduction in *R*^2^ when embeddings are compressed with methods other than mean pooling. A) Results for DMS data. B) Results for diverse protein sequences (PISCES data). In all cases, the y-axis represents different compression methods and the x-axis shows the resulting difference in *R*^2^. Dots represent the fixed effects estimates from mixed-effects modeling, and error bars represent 95% confidence intervals.

In aggregate, these results show that mean pooling is strictly superior in transfer-learning applications where the input sequences are widely diverged, and it performs well also with DMS data. Therefore, for our subsequent assessment of model size on transfer learning, we only considered mean pooling throughout.

### Moderately sized models perform well in transfer learning

We next turned to the effect of model size. We considered all six ESM-2 models, ranging from 8 million parameters (ESM-2 8M) to 15 billion parameters (ESM-2 15B). We also included the older model ESM-1v with 650 million parameters, which was developed specifically for variant effect prediction. Importantly, because ESM-1v only accepts sequences up to 1,022 residues, we excluded any DMS datasets with longer proteins from the main analysis, which reduced our DMS data to 36 distinct datasets. Results on the remaining datasets but excluding ESM-1v are presented in Supplementary Materials, Fig. S3. In addition to ESM-1v, we also evaluated several more recently published models, including AMPLIFY (120M, and 350M) and ESM C (300M and 600M, 6B). In aggregate, the models we considered range from small models with 8 million parameters to large models with 15 billion parameters. We categorized them into three groups based on parameter count: small (100 million), medium (100 million to 1 billion), and large (1 billion parameters). This classification is somewhat subjective, as it reflects the current state of available hardware. In a few years, billion-parameter models may be considered medium-sized. However, we chose this classification because of practical usability constraints in 2025—the large models according to our classification tend to require multiple high-end GPUs, limiting their accessibility.

We found that, overall, larger models tended to yield better results for transfer learning via feature extraction tasks across both types of datasets, as expected (Figure 3). However, the performance improvements for large models (with 3, 6, or 15 billion parameters) were moderate or small. For example, the medium-sized model ESM-2 150M consistently performed well across many targets—particularly in DMS analyses, where it slightly outperformed ESM-2 3B despite being twenty times smaller (Figure 3A). Additionally, all ESM C models outperformed ESM-2 15B, and scaling ESM C up to 6B offered only a minor advantage over the 600M parameter variant of the same model family. On the PISCES dataset, all models with 150M parameters or more delivered comparably strong performance, with the large ones reaching a close to optimal results (Figure 3B). These findings indicate that, for the datasets evaluated, the ESM C 600M model strikes an ideal balance between computational efficiency and predictive power, offering robust results in the context of transfer learning.

**Figure 3.**
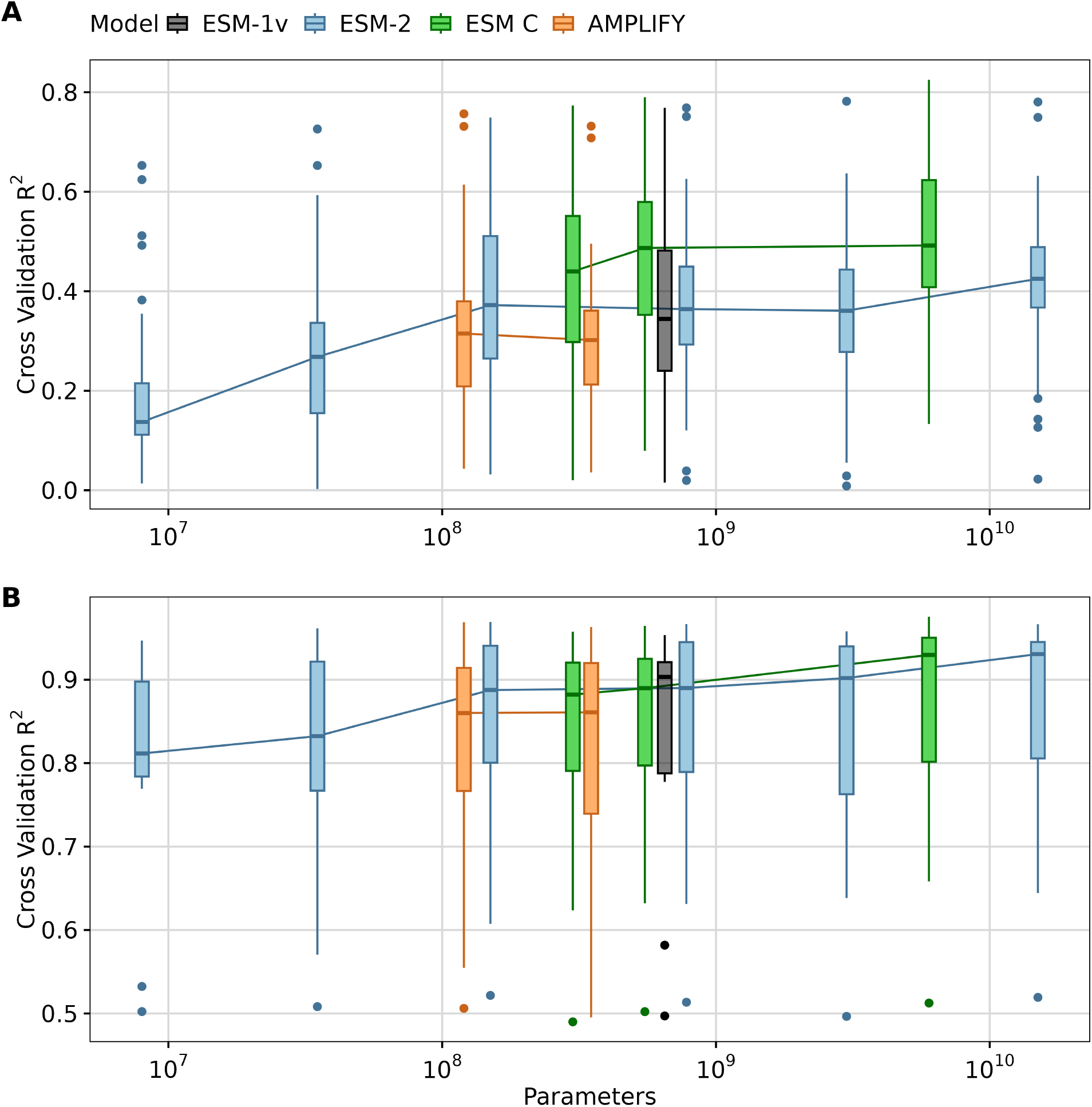
Impact of model size on transfer learning. A) LassoCV regression results using mean embeddings from different pLMs for 36 DMS datasets. B) LassoCV regression results using mean embeddings from different pLMs for 12 targets originated from the PISCES dataset. The y-axis displays the average *R*^2^ scores from 5-fold cross-validation for each dataset or target. The x-axis represents the number of parameters in each model, with different models grouped by color: ESM1v (black), ESM-2 models (blue), ESM C (green), and AMPLIFY (orange).

Although ESM-1v was specifically designed and trained on UniRef90^39^ for variant effect prediction, its limitation to sequences no longer than 1,022 residues restricts its applicability. To evaluate sequences exceeding this length, we turned to the ESM-2, ESM C, and AMPLIFY models. While these models were also pretrained with maximum sequence lengths of 1,024 for ESM-2 and 2,048 for ESM C and AMPLIFY, their use of a rotary attention mechanism allows them, in principle, to handle longer sequences. However, this capability remains underexplored. This part of our analysis was inconclusive, as none of the larger sequences (four in total) achieved particularly high predictability scores (Supplementary Materials, Fig. S3). Nonetheless, it is worth noting that ESM C models seem to display better performance overall for larger sequences, but more data is required to truly evaluate this question. At the very least, we can conclude that there is no strong evidence suggesting that ESM-2 and ESM C cannot be used with sequences longer than the maximum sequence length they were trained on.

To further investigate whether our conclusion—that larger models do not always deliver the best performance—holds for other transfer learning methods such as fine-tuning, we fine-tuned five ESM-2 models ranging from 8 million to 3 billion parameters, on 31 DMS datasets, resulting in 155 fine-tuned models. Due to significant computational resources required for fine-tuning pLMs, we had to restrict our experiment to DMS datasets with fewer than 15,000 samples and proteins shorter than 800 residues. We found that the fine-tuning results were qualitatively similar to the results with Lasso regression. While there was some improvement in performance with increasing model size, we saw a leveling off around 650 million parameters (Supplementary Materials, Fig. S4 and Fig. S5). In particular, fine-tuning a model 4.6 times larger (ESM-2 3B) did not outperform the medium-sized ESM-2 650M model. On the contrary, it sometimes showed a slightly worse performance. A similar observation was reported in a previous study^40^ when both models were fine-tuned for predicting protein–protein interactions (PPIs). In addition, we compared performance under fine-tuning to performance under feature extraction and found that performance was systematically higher under fine-tuning (Fig. S6).

### Sample size drives transfer learning performance in pLMs

Next, we investigated whether sample size impacts transfer learning performance. To explore this question, we selected three DMS datasets with good model performance (*R*^2^ > 0.6 for the larger models), large size ( 1000 observations), and differing complexity (single, double, and multiple mutations). We then progressively downsampled each dataset into subsets, ranging from 100 observations to the full dataset size. For each subset, we ran LassoCV regression with five-fold cross-validation. We found that smaller datasets negatively affected transfer learning via feature extraction performance, resulting in reduced accuracy for all three datasets and all three models (Figure 4). Notably, aside from the smallest two ESM-2 models (8M and 35M parameters) and AMPLIFY (120M and 350M parameter), all models performed comparably when sufficient data was available (≳ 10^4^ observations).

**Figure 4.**
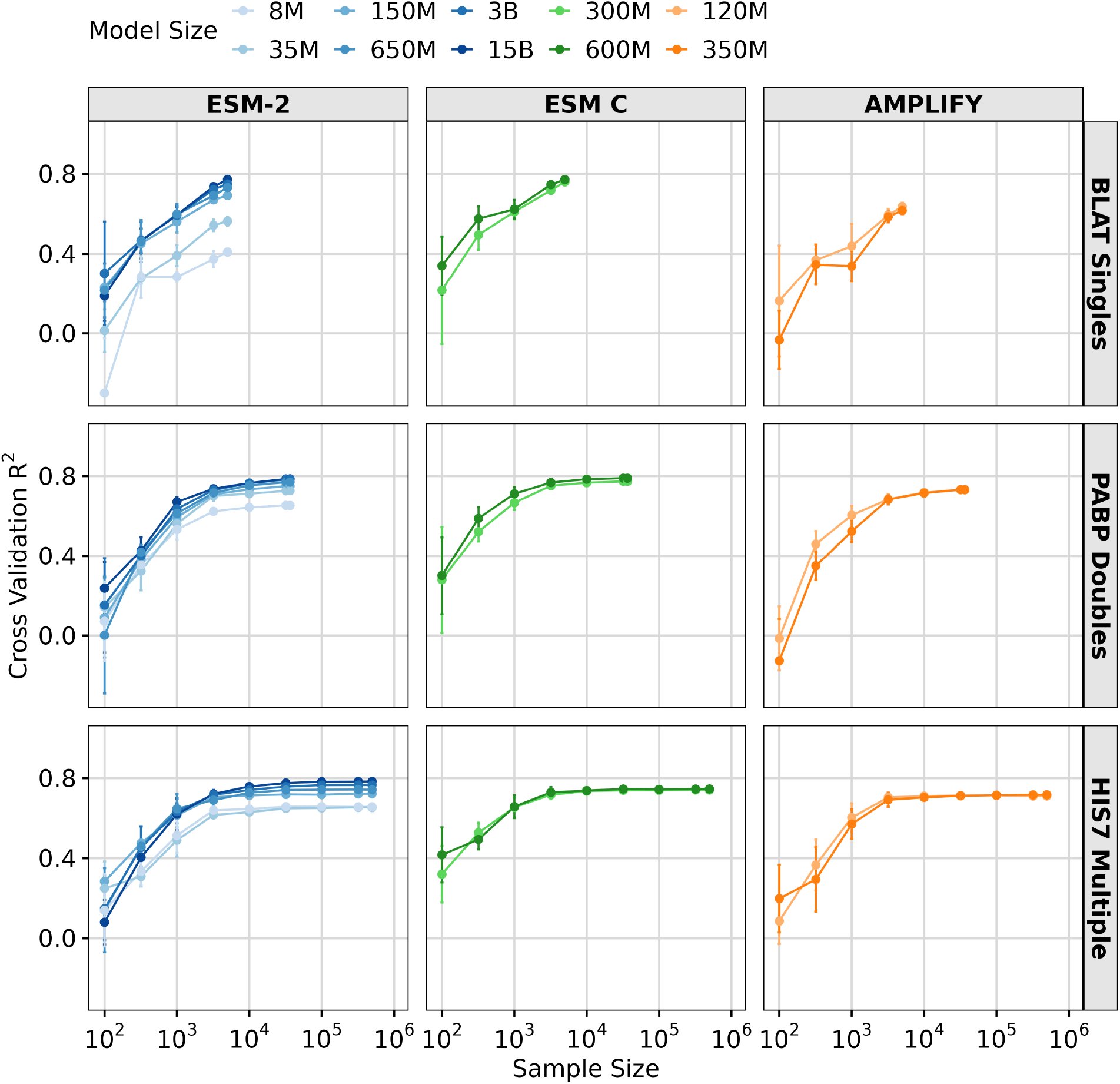
Effect of sample size on transfer learning via feature extraction. Results of LassoCV regression on three DMS datasets, using downsampled subsets ranging from 100 to the maximum number of samples in the dataset. The y-axis represents the averaged 5-fold cross-validation *R*^2^ scores, and error bars represent the standard deviation. The x-axis shows the sample sizes tested. The colored lines represent different models and model sizes: ESM-2 8M, 35M, 150M, 650M, 3B, and 15B (Blues), ESM C 300M, and 600M (Greens) and AMPLIFY 120M, and 350M (Oranges).

We hypothesized that meaningful protein features are distributed across more dimensions in larger models, leading to improved performance as the number of retained features increases. To test this hypothesis, we plotted the number of features retained by LassoCV for the three DMS datasets and models. We found that the ESM-2 15B model starts outperforming the smaller models when it starts utilizing a greater number of features (Supplementary Materials, Figure S7). Interestingly, the HIS7 dataset was the only one that showed clear saturation in the number of features being used across all models, for sample sizes in excess of ∼ 10^4^ (Supplementary Materials, Figure S7). For all models, the number of features used plateaued well below the models’ respective embedding dimensions. By contrast, in the other two datasets, for the largest models, the number of retained features continued to increase until the maximum dataset size was reached, suggesting that with a larger sample size performance could improve further. On the other hand, several medium-sized models showed this saturation with fewer features and a smaller sample size.

Interestingly, the number of features selected by LassoCV when using embeddings from ESM-2 650M and ESM C (300M and 600M) required only half of the number of features utilized by the larger ESM-2 15B model, and yet achieved comparable performance (Supplementary Materials, Figure S7). This result highlights the superior (or less sparse) protein representation encoded by the medium size pLMs, and in particular ESM C. Furthermore, utilizing fewer features offers significant benefits in transfer learning via feature extraction by reducing the risk of overfitting and by improving generalization. These results suggest that the larger embedding space provided by larger models cannot be fully taken advantage of by moderately sized datasets of fewer than 10^4^ observations, and this may be one of the main reasons moderately sized models are frequently sufficient and perform just as well as the largest models.

### Transfer learning is limited by data quality

Since the preceding analysis was performed by downsampling, where we expected prediction performance to systematically decline for more extreme ranges of downsampling, we next asked whether a similar relationship between dataset size and prediction accuracy could be observed across all DMS datasets, which inherently had differences in sample size. We additionally explored the effect of protein length and the type of data measured on model performance. We fit separate linear regression models, using transfer learning via feature extraction model performance (*R*^2^ score) as the dependent variable and either DMS dataset size or protein length as the independent variable. Our analysis revealed that the percent variation explained by sample size was minimal (for example, *R*^2^ = 0.004 for ESM C 600M, Figure 5A), indicating that while performance may improve with larger datasets, sample size is not the primary driver of transfer learning effectiveness in the 40 DMS datasets analyzed. Notably, most datasets had a sample size of over a thousand (Supplementary Materials, Fig. S8), a number close to optimal for transfer learning via feature extraction, as suggested by our downsampling results (Figure 4). However, a small sample size appears to have a more pronounced negative impact on the performance of the AMPLIFY family of models, which however also did not demonstrate strong overall performance (Supplementary Materials, Fig. S11).

**Figure 5.**
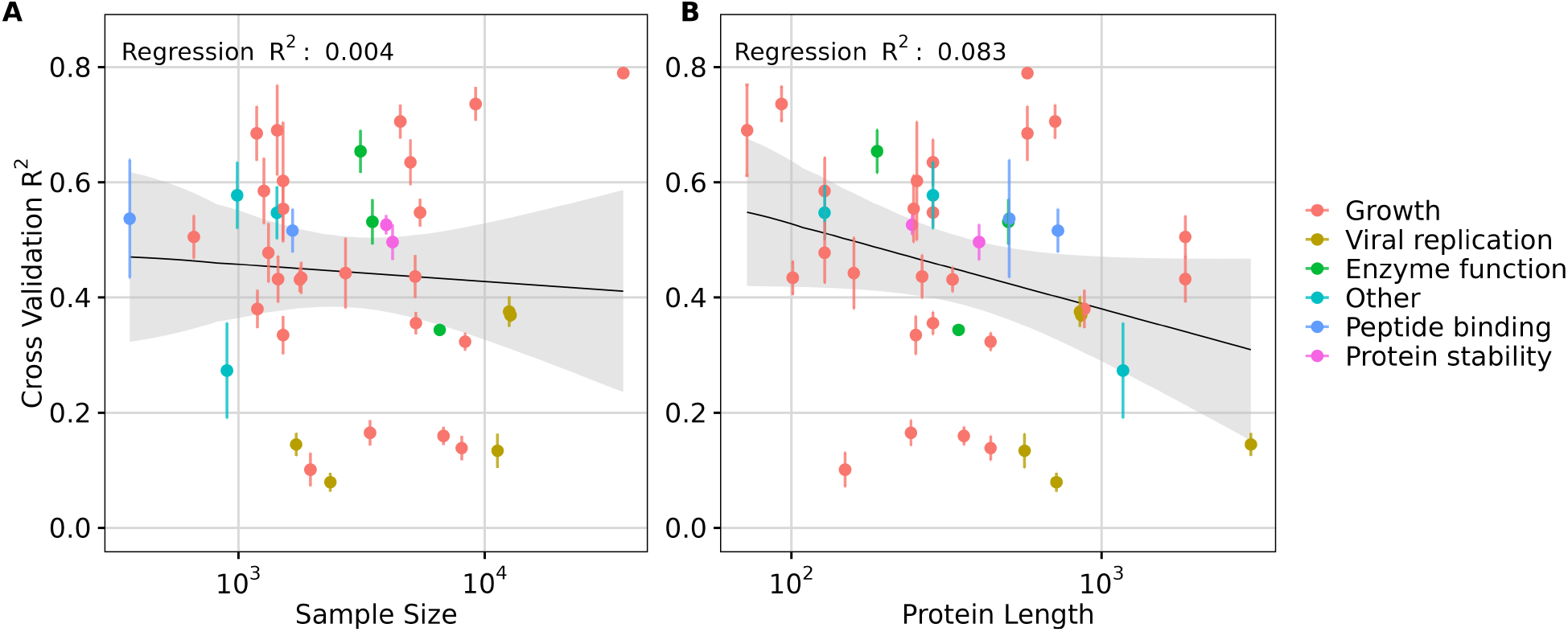
Effect of sample size and protein length in transfer learning via feature extraction. A) Effect of **sample size** on transfer learning using ESM C 600M model embeddings quantified by the regression results (*R*^2^ score). B) Effect of **protein length** on transfer learning using ESM C 600M model embeddings quantified by the regression results (*R*^2^ score). Each dot represents a dataset, and its color indicates the corresponding data measurement type. In each subplot, the y-axis represents the averaged 5-fold cross-validation *R*^2^ scores, and error bars represent the standard deviation.

By contrast, protein length accounted for a meaningful proportion of variation observed (for example, *R*^2^ = 0.083 for ESM C 600M), suggesting a modest influence of protein length on model performance. Specifically, longer protein sequences, particularly those exceeding the length of 1,022 residues, were associated with decreased model performance in transfer learning (Figure 5B and Supplementary Materials, Fig. S3). This trend was broadly consistent across different models (Supplementary Materials, Figs. S9, S10, and S11).

We also broke down the datasets by dataset type (growth, viral replication, peptide binding, etc.) but did not see a strong trend (Figure 5). One exception were viral proteins, which tended to be among the longest sequences in the DMS data and also exhibited some of the lowest *R*^2^ scores. This observation suggests that either pLMs embeddings may struggle to capture effective representations of viral proteins specifically, or that the embeddings generally lack representational power for larger proteins. We note that the vast majority of DMS datasets (all but four) used sequencing as the primary measurement method, and frequency measurements via sequencing are subject to numerous sources of noise^41^. These factors may partially explain the limited predictive power we observed for several of the datasets.

To investigate whether the variability in our results is influenced by the choice of embedding layer used for feature extraction, we performed a layer-wise analysis using the ESM-C 300M model. Prior work has shown that the final layer of a protein language model does not always yield the best performance in transfer learning^42^. To examine this effect, we extracted embeddings from all 30 layers for the 40 DMS datasets and evaluated their predictive performance using LassoCV regression. This analysis revealed that performance tended to increase in deeper layers of the neural network (Figure S12). Across all datasets considered, the last layer was most commonly the top-performing layer, and in cases where it was not it typically the penultimate layer performed best (Figure S13), with the final layer having just slightly lower performance (Figure S12). Based on this analysis, we concluded that the last layer was the most effective and straightforward choice for our analysis and was not the main factor behind the low predictive performance observed in some datasets.

## Discussion

We have evaluated the performance of the ESM-2, ESM-1v, ESM C, and AMPLIFY model families across various model sizes (from 8 million to 15 billion parameters) in transfer learning tasks on a wide range of different biological datasets. We have found that larger models do not consistently outperform smaller ones, especially when data is limited. In fact, medium-size models, such as ESM-2 150M, ESM-2 650M or ESM C 600M, have demonstrated consistently good performance. This observation suggests that model size should be carefully aligned with dataset size and data type to optimize transfer learning performance. Additionally, we have evaluated different methods for compressing per-token embedding matrices, and we have found that, on average, mean embeddings consistently outperform other commonly used compression methods. However, for fairly homogeneous DMS datasets, alternative compression methods including max pooling and iDCT can on occasion achieve better performance than mean embeddings. Finally, we have found that fine-tuning tends to outperform transfer learning via feature extraction, though the results of the two methods are correlated. In particular, our conclusions about model size appears to apply equally to both learning approaches.

### Medium-size models perform well in transfer learning

Scaling large language models (LLMs) to ever larger numbers of parameters has become a trend since Kaplan et al.^43^ showed that there is a power-law relationship between model size and performance. This scaling law has driven the development of increasingly larger models^26,27^. Since the introduction of Generative Pre-Trained Transformers (GPTs)^44^ in 2018, LLMs have scaled to trillions of parameters^45^. This tendency has also extended to biology, with the latest protein language models (pLMs) reaching an astonishing 98 billion parameters^7^. Although larger pLMs have demonstrated enhanced performance by capturing more complex relationships in protein sequences^16^, a recent study has shown that as models scale, they become more prone to overfitting, often preferring to predict the wild-type residue in masked tasks^29^. This observation aligns with the known LLM scaling laws, which suggest that performance improvements from scaling are only achieved if the amount of training data is scaled as well^43^. As ESM-2 models were not trained with data scaled across different model sizes, their performance gains may have been limited, in particular for the larger models.

Smaller models offer many advantages, such as reduced inference costs and improved compatibility with compact hardware, providing substantial benefits to protein studies. In the context of regular (i.e., non-biological) large language models, Hoffmann et al.^46^ showed that training a smaller model with more data (70 billion parameters and 1.4 trillion tokens) outperformed larger models with four times more parameters when pre-trained on less data. Our findings show similarly that the more recently developed smaller ESM C models in many cases outperform the older ESM-2 models, even for the model variants that have over an order of magnitude more parameters. While the model architecture of ESM C models is slightly different from ESM-2 models, we suspect the main benefit of the newer models comes from an improved training schedule. Overall, our results align with other studies indicating that pLMs beyond 650 million parameters have limited advantages in many application areas, only showing true benefits when used for tasks focused on structure prediction^47–49^.

While our study has primarily focused on quantitative comparisons across model sizes and dataset types, several qualitative patterns have emerged that provide insight into observed performance differences across datasets. Notably, viral protein datasets tend to yield lower predictive performance. These proteins are often longer than those in other datasets, and our observations suggest that all pLMs we considered struggle to capture meaningful representations for longer sequences. This could be due to limitations in the model’s effective context length, which is typically about half of the maximum sequence length the model was trained on^50^, or maybe due to a lower representation of viral proteins in the training data. Additionally, we have observed that the majority of DMS datasets rely on sequencing-based measurements, which are prone to experimental noise, such as PCR amplification errors^51^. Such noise can obscure the underlying biological signal, hindering machine learning model predictions^52^. These observations highlight that both biological and experimental factors—not just model architecture or size—play a critical role in model performance.

### Impact of embeddings compression strategies and layer selection

In addition to evaluating model size, we also systematically evaluated various feature compression methods. Compression methods are required to reduce the position-dependent embedding matrix generated by pLMs into a comparatively much lower-dimensional feature vector whose dimension is independent of the protein length. The compression method iDCT in particular has been proposed as a better alternative to mean pooling^24^. However, we could not recapitulate this finding here, except for some specific DMS datasets. As a general and simple approach, mean pooling seems to be the best option. We caution however that our results do not imply no better compression methods exist. For any particular dataset, it is generally possible to find a better dataset-specific compression method, by training the compression method on the data^53–55^. Our recommendation here is to use mean pooling as a reasonable starting point, and then potentially consider dataset-specific compression methods in case mean pooling does not yield adequate results.

Along the same lines, we used the final embedding layer of each model for transfer learning but other choices are possible. In particular, there is some evidence in the literature that using intermediate layers or multiple layers can yield better transfer learning performance than using the last layer^24,48,56^. However, all available evidence indicates that any optimizations with regards to optimal layer choice will be dataset dependent. For any specific application task, it may always be worthwhile to perform optimizations with respect to which pLM, compression method, or embedding layer(s) are used, as long as good care is taken that a final test dataset is set aside and has never influenced any of these modeling choices. In fact, the comprehensive prediction benchmark ProteinGym has previously suggested that model selection should be guided by the dataset itself, as performance can vary significantly across assays^57^.

### Reproducibility challenges with pLMs

We note that while working on this project, we encountered numerous reproducibility issues with computed embeddings that we think the field needs to be aware of. These issues arise because calculating embeddings for even a single protein sequence requires billions of floating-point operations and small rounding errors or other inaccuracies can accumulate to the point where the final result has widely diverged from what it should be. The only strategy that avoids this issue entirely is to calculate embeddings on the exact same hardware architecture and GPU drivers that were used to train a given model. In practice, this will frequently not be possible. However, we found that models can be re-used across architectures, even from different vendors (NVIDIA vs AMD vs Apple Silicon), as long as the user is aware of the most common pitfalls and ensures they are avoided.

We can group reproducibility issues into three distinct categories: (i) issues with software libraries/drivers; (ii) issues due to numerical types; and (iii) issues due to batch processing of multiple sequences at once. We encountered the first issue when trying to run ESM-2 models on Apple Silicon GPUs via MPS (Metal Performance Shaders). We found that calculated embeddings had accumulated rounding errors to the point that even the first significant digit in most embedding dimensions differed from the correct value. Issues filed on the PyTorch GitHub repository (e.g., issue #84936) suggest that these problems were due to the specific floating point math implementation used by Apple in their drivers. We note that these issues seem to have been resolved since the release of MacOS Sequoia and PyTorch version 2.5 in the fall of 2024 but may still be present on many Apple devices not yet updated to the latest software.

We encountered problems with numerical types in the ESM C models, which by default use reduced accuracy floating point types and autocasting to speed up performance during inference. Autocasting in particular dynamically alters numerical precision, potentially leading to inconsistencies across runs or hardware configurations. These precision variations were more prominent when passing an input array with multiple sequences, and they were overall sufficiently severe that we felt we could not generate reproducible embeddings from ESM C models in the default configuration. However, we were able to circumvent these issues by using the `float32` type consistently throughout and disabling autocasting.

Finally, we encountered issues with padding with the AMPLIFY models. In all models, when processing a batch of multiple sequences with different lengths, shorter sequences need to be padded to the maximum length. This padding should not affect computed embeddings, but if a transformer model does not properly mask padded sites when calculating attention then the padding can influence output embeddings, which will result in poor reproducibility. In models with this issue, the embeddings for one sequence will depend on the lengths of other sequences processed in the same batch. We were able to circumvent this issue by not using any batch processing and always processing each sequence individually without any padding.

Fortunately, we have found that it is relatively straightforward to check for the presence of any of these reproducibility issues. Embeddings calculated on a CPU using the `float32` type, on individual sequences without batching, seem to always be reliable, for all models. Therefore, we recommend to always compare GPU results to this gold standard and to only proceed if any observed differences between CPU and GPU results are minor.

### Limitations

Our study has several limitations. First, we have focused on ESM-2 and related models (ESM-1v, ESM C, AMPLIFY). ESM-2 has the widest range of model sizes and therefore is a logical starting point for an analysis on the impact of model size. To keep the study manageable, we excluded other models, such as the ProtTrans family^10^ or ProtFlash^11^. We also excluded generative models such as ProGen^12^. We do not believe that including these models would have substantively altered our conclusions, in particular since most of these models appear to perform substantially worse than ESM C or even ESM-2^30^.

Second, our analysis of embedding compression methods considered relatively simple approaches, such as mean pooling, max pooling, extracting just the embedding corresponding to the beginning-of-sequence (BOS) token, and various dimension reduction techniques such as PCA and iDCT. Among those, mean pooling was consistently outperforming the other methods. However, we acknowledge that methods that train the compression method on the data^53–55^ may outperform mean pooling.

Third, we primarily focused on transfer learning via feature extraction, using the final embedding layer. Other layers may perform better, though this was not typically the case in our DMS data. More importantly, fine-tuning may yield higher performance than LassoCV on extracted features. We did not systematically explore fine-tuning due to the substantially higher compute required to do so. To the extent that we did do fine-tuning, we observed that results were broadly correlated with results from LassoCV, suggesting that our overall conclusions are not dependent on the specific transfer learning method (feature extraction versus fine-tuning).

Finally, we would like to emphasize that despite the marginal performance gains observed in our analysis with the largest pLMs, with 3 billion or more parameters, their practical use presents several limitations. Most importantly, these models require substantial computational resources. For example, we were unable to fine-tune ESM-2 15B even with high-memory GPUs such as the NVidia H100 and using parameter-efficient methods such as LoRA. And even for the smaller ESM-2 3B model we were unable to fine-tune with the largest datasets and datasets containing the longest proteins. Furthermore, large models typically require large datasets to fine-tune effectively. In many biological applications, experimental data is limited, which constrains the ability to adapt these models to specific tasks. While it is possible that models like ESM-2 15B could yield better results under ideal conditions, these technical and resource constraints made them largely inaccessible for our experiments.

## Conclusion

In summary, our study challenges the general preference for larger protein language models in biology. In particular, it suggests that in common transfer learning applications, most available datasets are too small for the largest available pLMs. While larger models can achieve good performance in principle, medium-size models frequently perform similarly or deliver stable performance with smaller samples, making them more accessible and practical in the more common scenarios working biologists may experience in practice. Additionally, mean embeddings consistently perform well across datasets, suggesting that alternative compression methods will rarely be needed. In general, mean embeddings calculated from models of intermediate size, such as ESM C 600M, are likely sufficient for transfer learning on the vast majority of data sets encountered in biological research.

## Methods

### Data collection

First, we obtained 41 deep mutational scanning (DMS) datasets previously curated in Ref.^31^. A complete list of all datasets included is available at: https://github.com/ziul-bio/SWAT/blob/main/data/DMS_metadata.csv. Based on the collected data, we created FASTA and CSV files containing the mutated sequences for each protein, along with the corresponding target values of the mutations. The code to reconstruct sequences is available at: https://github.com/ziul-bio/SWAT/blob/main/notebooks/DMS_pre_processing.ipynb.

Second, we downloaded the PISCES dataset^32^, which contains a collection of proteins with known structures and at most 50% pairwise sequence similarity. We filtered this dataset to include only proteins with lengths between 64 and 1022 residues, resulting in a total of 23,487 sequences. The minimum length of 64 residues was determined by a requirement in our implementation of the inverse Discrete Cosine Transform (iDCT, see below), while the maximum length of 1022 residues corresponds to the upper limit supported by ESM-2 models. The final dataset is available at: https://github.com/ziul-bio/SWAT/blob/main/data/PISCES/pisces_len64-1022.fasta. From this dataset, we generated 12 target variables, including a selection of physicochemical properties, amino acid frequencies, and secondary structure frequencies (Supplementary Materials, Table S1). For physicochemical properties, we selected six interpretable features from those available in the Peptides python package (version 0.3.2), accessible on PyPI. For the selection of amino acid frequencies, we arbitrarily chose alanine, cysteine, and leucine. For secondary structure targets, we selected the three most common ones (alpha helix, beta strand, and coil), since all others were rare in most sequences. Secondary structure targets were computed using the DSSP module provided by Biopython. The code required to generate these targets is available at: https://github.com/ziul-bio/SWAT/blob/main/notebooks/PISCES_pre_processing.ipynb

### Computing protein embeddings from ESM-2 and ESM1v

We calculated protein embeddings from the ESM-2 family of protein language models^16^ and from the older ESM-1v model^58^ using the same approach. For all model variants, the model’s internal representation is a matrix of embeddings in **R**^*n*×*d*^ dimensions, where *n* represents the protein sequence length and *d* represents the embedding dimension, which differs for different model variants and generally increases for models with more parameters. For each protein sequence in our datasets, we fed the sequence into the respective model and obtained the corresponding embeddings from the final hidden layer. For all models, we obtained embeddings using the `extract.py` script available at the ESM-2 GitHub repository (https://github.com/facebookresearch/esm/blob/main/scripts/extract.py). This script allows us to define three output representations: mean representation (the embeddings averaged across the sequence length), BOS representation (the CLS token or beginning of the sequence), and per token representation (the full embedding matrix). We used this script to obtain all three representations. The mean and BOS representations were used as it is, and the per token representation was further processed as discussed below.

To extract ESM-2 embeddings of sequences longer than 1022 residues, we modified the script by increasing the max truncation length of sequences to 5000. See the script available at: https://github.com/ziul-bio/SWAT/blob/main/scripts/extract_esm2.py

### Computing protein embeddings from ESM C

To extract protein mean embeddings from the ESM C models, we followed the instructions provided on the Evolutionary Scale GitHub page: https://github.com/evolutionaryscale/esm. Based on these instructions, we developed a custom extraction script, available at: https://github.com/ziul-bio/SWAT/blob/main/scripts/extract_ESMC.py. Before running any of the ESM C models, we had to make a few modifications to ensure the reproducibility of results. First, we changed the data type from `bfloat16` to `float32` in the `esmc.py` file. The `bfloat16` data type is optimized for efficiency and reduced memory usage during training, but due to its half-precision nature, it can sometimes introduce numerical precision issues. By switching to `float32`, we ensured higher precision, which is particularly important when generating embeddings for downstream tasks or analyses. This adjustment also ensured consistency in results independent of the hardware being used. Second, we disabled autocasting, by setting `enabled=FALSE` and `dtype=torch.bfloat32` in the `torch.autocast()` function. Autocasting dynamically adjusts the precision during computations to optimize performance, but this can lead to inconsistencies in results across different runs or hardware setups. Disabling autocasting and explicitly setting the data type to `float32` ensures that all calculations are performed with consistent precision, making the embeddings more reliable and reproducible.

As of this writing, the ESM C 6B model is not publicly available. Instead, access requires enrollment in the beta access program and the creation of an access token for API calls. Due to this limited access, we were unable to obtain results for the DMS HIS dataset, which contains 500,000 sequences. Another limitation concerns the maximum sequence length: While ESM C 300M and 600M can handle sequences longer than 1022 residues out of the box without any modifications, ESM C 6B, accessed through the API, has a preset maximum sequence length of 2048. This restriction excluded the dataset ‘POLG Sun2014’ which the protein has 3033 residues.

### Computing protein embeddings from AMPLIFY

To extract embeddings from the AMPLIFY model, we built a custom Python script, available at: https://github.com/ziul-bio/SWAT/blob/main/scripts/extract_AMPLIFY.py. We then downloaded the model through Hugging Face, and after having the model locally, we made a few modifications to ensure reproducibility. First, we normalized the last hidden layer embeddings. The ESM-2 and ESM C models have the output of the last hidden layer normalized with `pytorch.nn.LayerNorm`. To ensure that results were comparable between these models and the AMPLIFY model we did the same for the AMPLIFY embeddings, which are not normalized by default. Second, AMPLIFY has a maximum sequence truncation defined as 2048. To be able to handle longer sequences, we modified the file `config.json` downloaded with the model and changed the parameter `max_length` to 5000.

### Per-token embeddings compression

To compress the per-token representation matrix, we employed several techniques from scikit-learn^59^, such as `MinMaxScaler`, PCA (Principal Component Analysis), and Kernel PCA. Initially, the embeddings matrix was scaled using `MinMaxScaler(feature_range=(-1, 1))` applied separately to each feature dimension along the protein sequence. To compress the embeddings, we applied PCA to the scaled embeddings matrix, treating the sequence length as features and the model dimensions as samples and retaining the first two PCs. Similarly, we used kernel PCA with either radial basis functions (RBF) or sigmoid kernels, using the scikit lern function `KernelPCA()`. Following the transformations to extract a single vector of fixed length, the first and second components of the PCAs were used and subsequently named PC1/PC2, RBF PC1/RBF PC2, and Sigmoid PC1/Sigmoid PC2, respectively.

Lastly, we employed an inverse Discrete Cosine Transform (iDCT) quantization method as a method of embeddings compression, as described in the Protein Ortholog Search Tool (PROST)^24^. iDCT is particularly adept at capturing fine-grained details in data while reducing dimensionality by discarding high-frequency components, enhancing the signal-to-noise ratio in the embeddings^60,61^. We applied the Discrete Cosine Transform (DCT) to the embeddings, retained the top rows and columns, and then performed the inverse DCT (iDCT) to reconstruct the embeddings. The reconstructed matrix was reshaped into a 1D vector, with final length determined by the number of retained rows and columns. We experimented with various configurations, retaining subsets of rows and columns such as 5x44, 10x64, 10x128, 10x512, and 10x640. These configurations correspond to fixed-length embeddings of 220, 640, 1280, 5120, and 6400, respectively, which we designated as iDCT1, iDCT2, iDCT3, iDCT4, and iDCT5.

### Statistical Analysis

To assess the statistical significance of differences among compression methods, we employed linear mixed-effects models using the lme4 package in R^62^. This method is well-suited for analyzing data with hierarchical or nested structures. In our analysis, the fixed effect was the compression method (e.g., Mean, BOS, PCA, kernel PCA, and iDCT), while the random effect accounted for variability across the different datasets. To evaluate the significance of the fixed effects estimates we used the package multcomp^63^ in R to calculate 95% confidence intervals around each estimate, taking into account that we were testing multiple hypotheses at once (one for each compression method).

### Predicting Protein Fitness and Properties

To evaluate the predictive performance of compressed protein embeddings, we employed LassoCV, a cross-validation-based version of the Lasso algorithm^64^. LassoCV retains the feature selection properties of Lasso by reducing the coefficients of less relevant features to zero while selecting the optimal regularization parameter through cross-validation, thereby identifying the most important predictors.

All modeling was conducted in Python using methods from scikit-learn, including LassoCV, cross-validation (KFold), and performance metrics (*R*^2^, MAE, RMSE, and Spearman’s *ρ*). Before any modeling, the dataset features were scaled using `MinMaxScaler(feature_range=(0, 1))`. Using a 5-fold cross-validation strategy, the data was then divided into five parts, with four parts used for training and one for testing during each round of cross-validation.

To determine the optimal regularization parameter *α* within each round of cross-validation, the LassoCV algorithm as implemented by scikit-learn generates a sequence of 100 *α* values, spanning three orders of magnitude in range. For each *α*, Lasso regression is performed on the training data, and a 3-fold cross-validation evaluates model performance. The optimal *α* is selected based on the lowest cross-validation error, and a final Lasso model is fit using the best *α* and the entire training dataset. Using this final Lasso model, we then calculated performance metrics for both the training and the test set.

This process was repeated five times and the average and standard deviation of the performance metrics on the cross-validation test sets are reported.

### Fine-tuning ESM-2 models

To fine-tune the five ESM-2 models (ranging from 8 million to 3 billion parameters) presented in our work, we leveraged PFET (Parameter-Efficient Fine-Tuning)^65^ to apply LoRA (Low-Rank Adaptation)^66^ layers (specifically, `q_proj` and `v_proj`) to the models while freezing all other layers. The LoRA layers were set with a rank of 4, a scaling factor of 32 (`lora_alpha`), and a dropout rate of 0.01. As a modification to the original model architectures, we replaced the language modeling head with a regression head. The regression layer receives as input the model’s embedding vector corresponding to the BOS (beginning of sequence/classification) token. For each of the ESM-2 models, the regression head began with a linear projection from the embedding vector to a set of hidden nodes matching the embedding dimension of the model being fine-tuned, followed by dropout (rate 0.1), layer normalization, and an output projection to produce the final regression value.

We selected 31 DMS datasets for fine-tuning, using only those datasets with fewer than 15,000 samples and proteins shorter than 800 residues. Each dataset was then randomly split into training and test sets (at a 80/20 ratio) using the same random seed to ensure consistent splits across all fine-tuning runs. To fine-tune, models were trained for 40 epochs with a batch size of 8.

The learning rate was set to 1 × 10^−4^ with a weight decay of 1 × 10^−6^. Due to the high computational cost of fine-tuning 155 models (31 datasets times five model sizes), each model was fine-tuned only once per dataset. The best model was chosen based on the epoch immediately following the intersection of the validation and training *R*^2^ values, ensuring that the training *R*^2^ was not more than 0.06 higher than the validation *R*^2^, which was the mean difference observed in our LassoCV results. Training was performed on AMD Instinct™ MI100 GPUs using PyTorch^67^, and the final outputs were saved to CSV files.

### Layer-wise analysis in protein language model

To evaluate the contribution of each layer within the protein language model, we conducted a systematic analysis of all layers in the ESM-2 300M model. Specifically, we extracted mean embeddings from all 30 layers across 40 deep mutational scanning (DMS) datasets, as detailed in the subsection “Computing Protein Embeddings from ESM-2”. Following this, we predicted protein fitness using LassoCV, as described earlier, for each of the 30 layers, performing this procedure independently for each DMS dataset.

## Code availability

All code used for data processing, embedding extraction and compression, regression models, fine-tuning scripts, and plotting results is available at: https://github.com/ziul-bio/SWAT.

## Acknowledgments

This work was supported in part by National Institutes of Health grants R01 AI169462, R01 AI148419, R01 GM088344, and R56 AI179799. The Texas Advanced Computing Center (TACC) provided high-performance computing support. C.O.W. also acknowledges support from the Blumberg Centennial Professorship in Molecular Evolution. The authors thank the Biomedical Research Computing Facility at UT Austin, Center for Biomedical Research Support (RRID: SCR_021979), for providing the computational resources used to fine-tune the pLMs. We also thank Aaron Feller for support and discussions on the topic of large language models.

## Author contributions statement

L.C.V. designed the study, performed data analysis, interpreted results, and wrote the manuscript. M.L.H. assisted with computational implementation, data preprocessing, and visualization. C.O.W. provided supervision, conceptual guidance, and critical manuscript revisions. All authors contributed to manuscript writing and approved the final version.

## Additional information

