## Supplementary materials for "Medium-sized protein language models perform well at transfer learning on realistic datasets"

May 8, 2025

### Supporting Information

#### Supporting Figures

#### Supporting Table

*All authors affiliations are: Department of Integrative Biology, The University of Texas at Austin, Austin, TX, United States of America

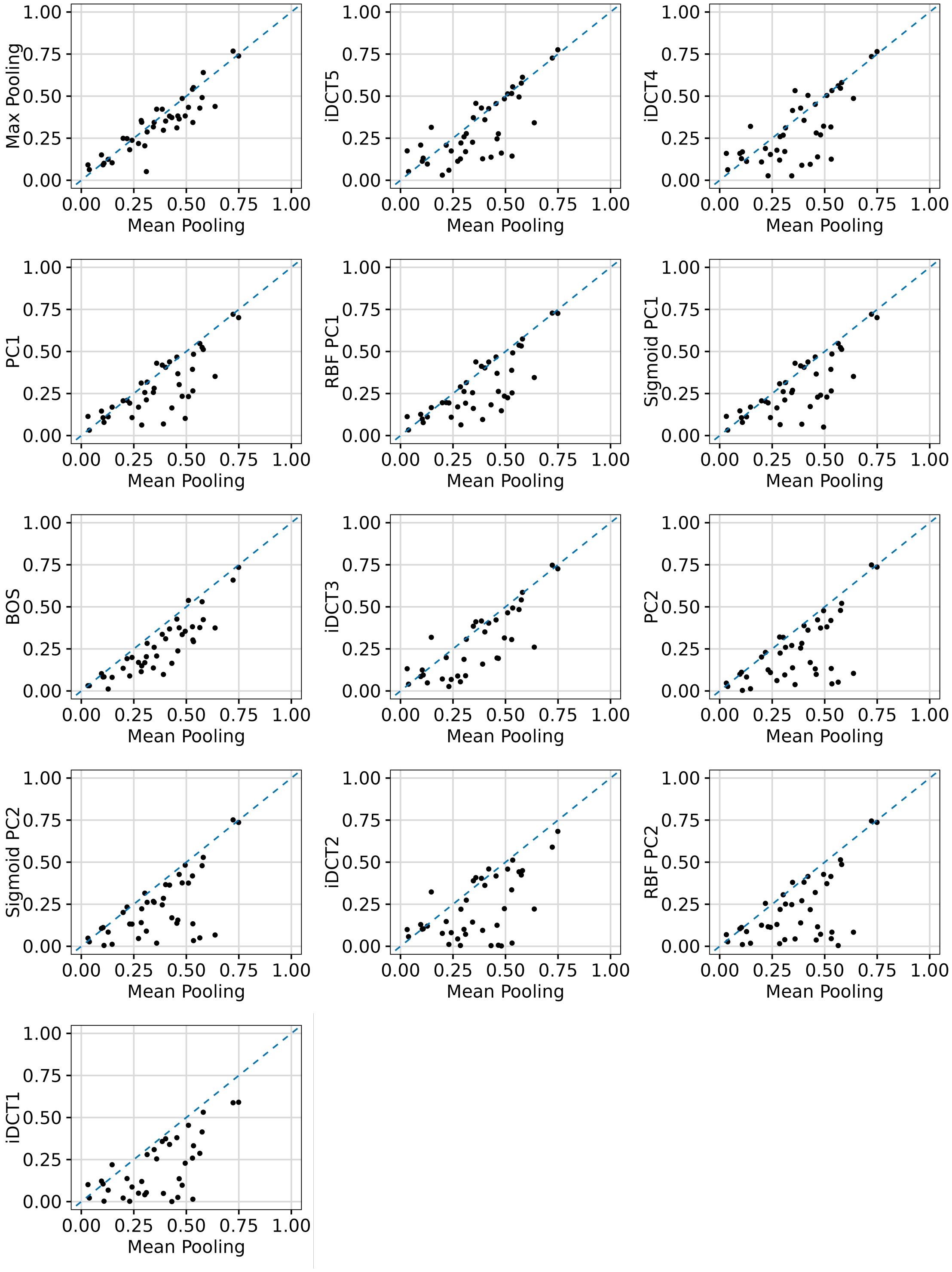


Figure S1: **Comparison of mean embeddings with 13 alternative compression methods on DMS datasets.** In each plot, the x-axis represents the averaged $R^2$ scores from 5-fold cross-validation for the mean embeddings, while the y-axis shows the averaged $R^2$ scores from 5-fold cross-validation for one of the other 13 compression methods. Each dot corresponds to one of the 40 DMS datasets analyzed.

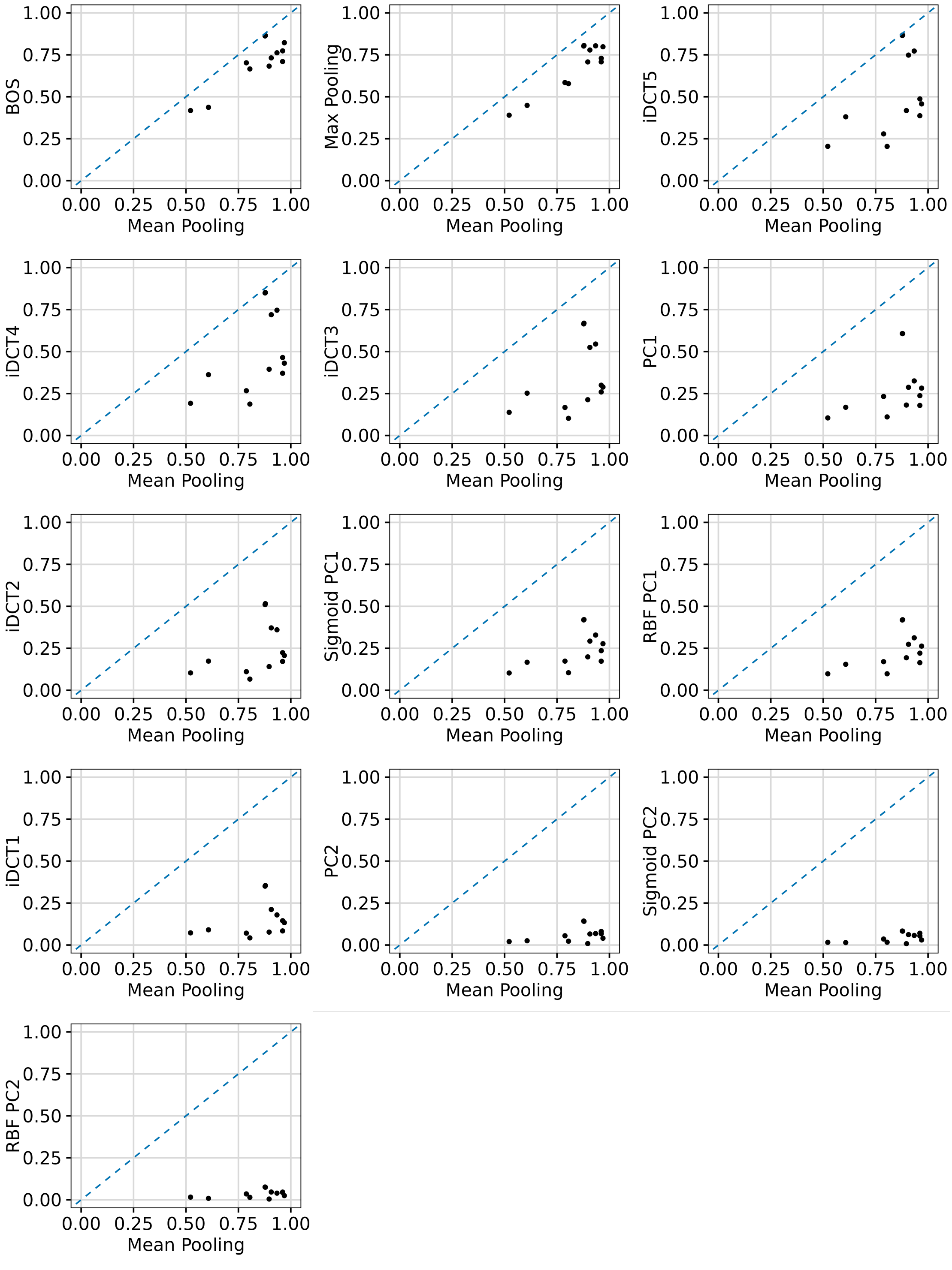


Figure S2: **Comparison of mean embeddings with 13 alternative compression methods on PISCES dataset.** In each plot, the x-axis represents the averaged $R^2$ scores from 5-fold cross-validation for the mean embeddings, while the y-axis shows the averaged $R^2$ scores from 5-fold cross-validation for one of the other 13 compression methods. Each dot corresponds to one of the 12 targets analyzed from PISCES dataset (PCPs and Secondary Structure).

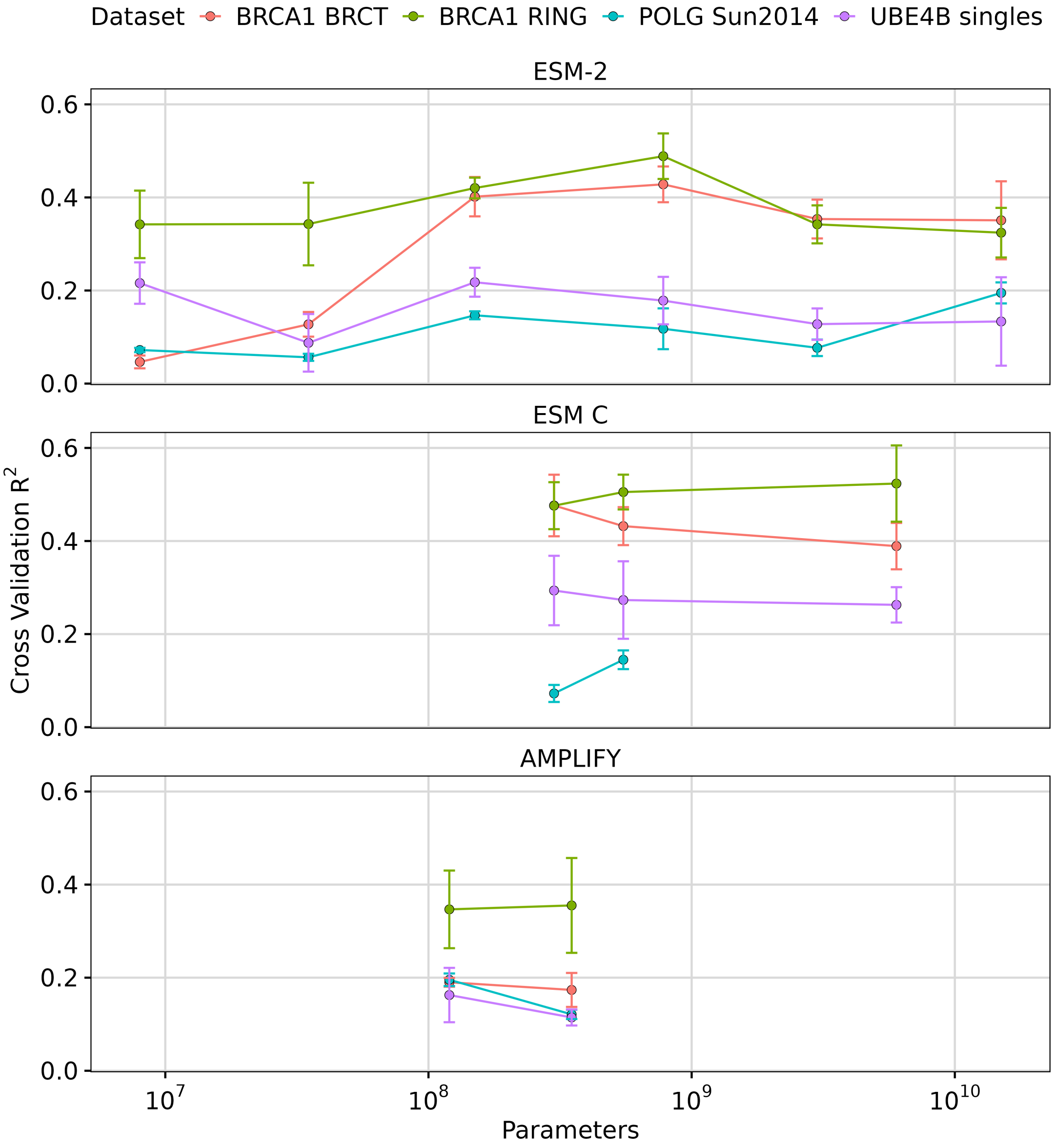


Figure S3: **Performance of pLMs on larger protein sequences.** Prediction scores across 4 different DMS datasets with sequence lengths greater than 1022. The y-axis represents the averaged 5-fold cross-validation $R^2$ scores, and error bars represent the standard deviation. The x-axis shows the different model sizes. Colors represent individual protein datasets, varying in their lengths: 3033 (POLG Sun2014), 1863 (BRCA1 BRCT), 1863 (BRCA1 RING), 1173 (UBE4B singles).

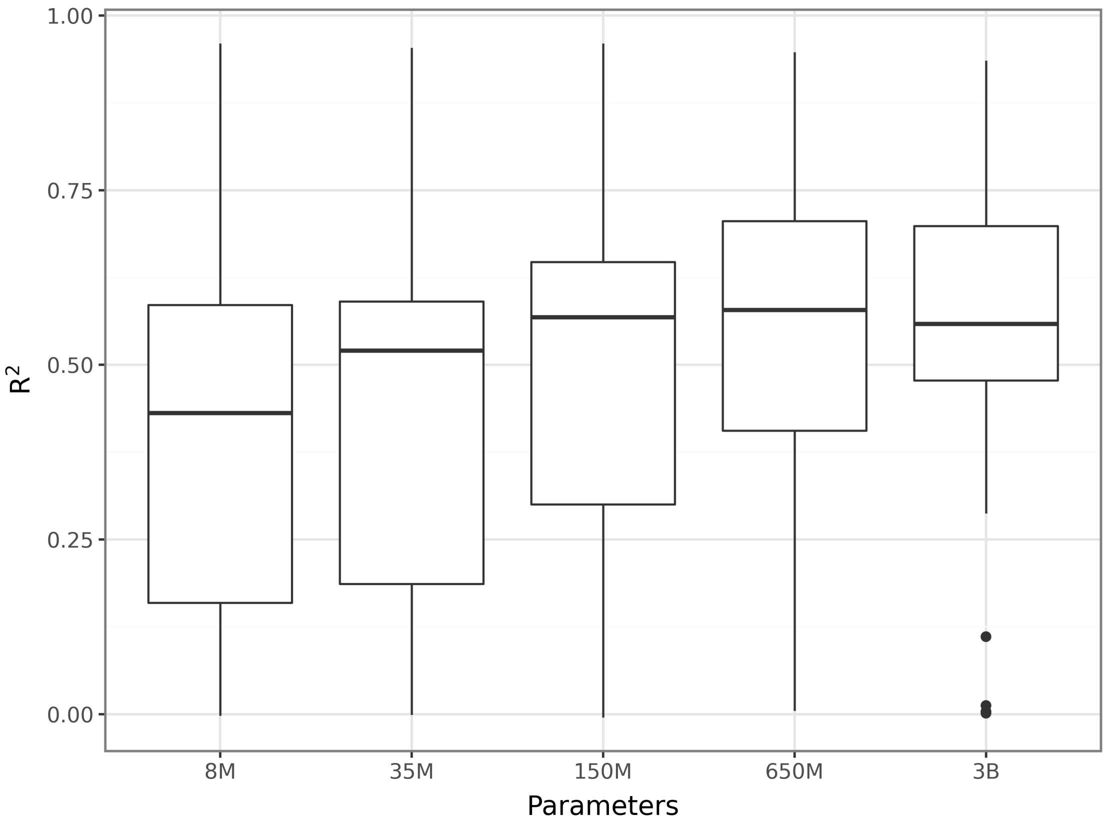


Figure S4: **Impact of model size on fine-tuning ESM-2.** We are plotting the distribution of $R^2$ scores obtained for 31 different DMS datasets against the number of parameters in the respective ESM-2 model.

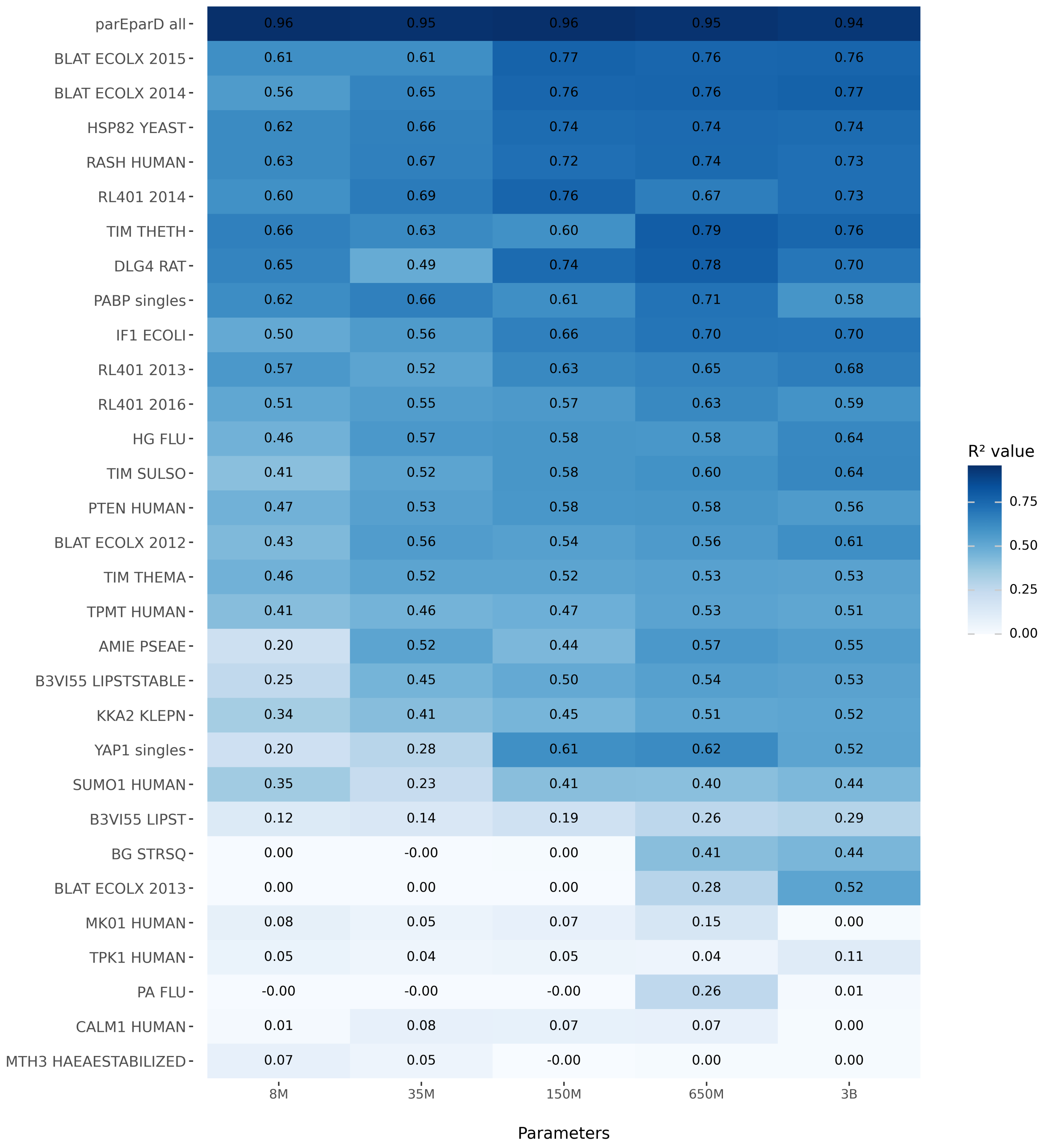


Figure S5: **Impact of model size on fine-tuning ESM-2 per dataset.** We plot the individual $R^2$ scores obtained for 31 different DMS datasets against the number of parameters in the corresponding ESM-2 models. The $R^2$ scores are shown in blue, with their corresponding values annotated. Rows are sorted by the average $R^2$ across models for each dataset.

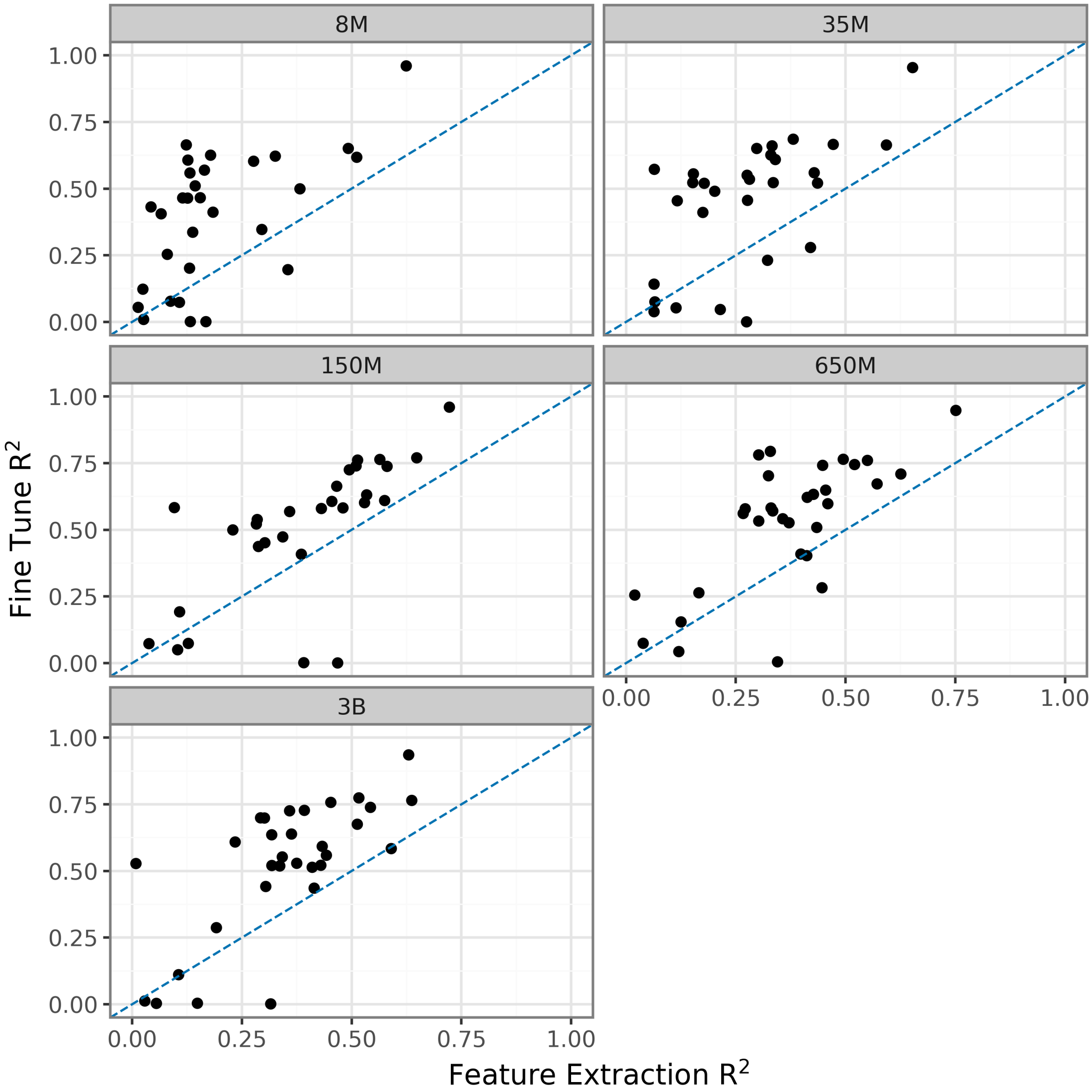


Figure S6: **Comparison between transfer learning strategies.** We are plotting the $R^2$ scores obtained by fine-tuning ESM-2 versus the $R^2$ scores obtained by transfer learning via feature extraction and LassoCV, for the same 31 DMS datasets as in Figure S4. The panels correspond to the different ESM-2 model sizes.

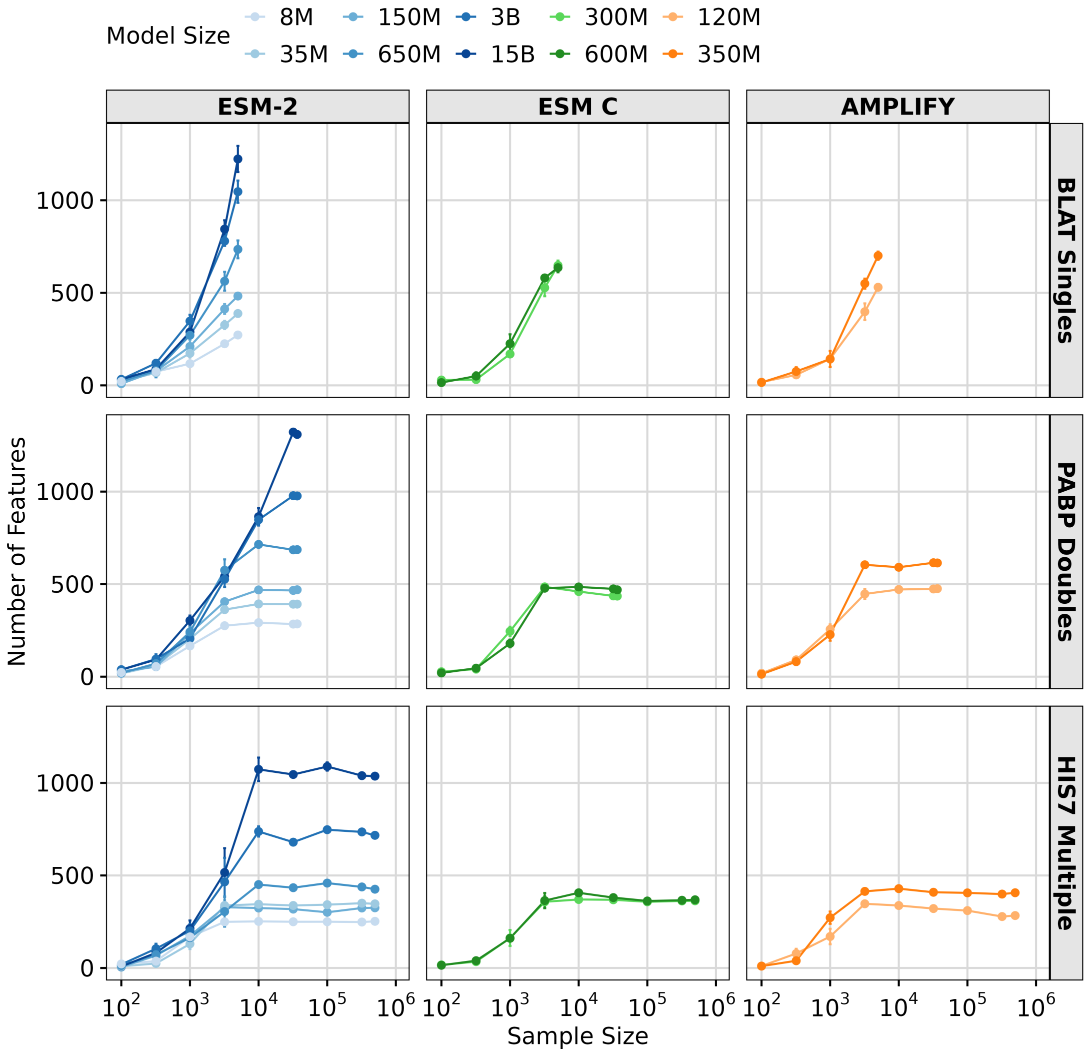


Figure S7: **Number of features boosts performance in protein prediction tasks.** Number of important features identified by LassoCV regression for three DMS datasets with varying mutation complexities: single mutations (BLAT ECOLX - 2015), double mutations (PABP), and multiple mutations (HIS7 YEAST). The y-axis represents the averaged 5-fold cross-validation number of important features (or non-zero coefficients set by Lasso l1 penalization) scores, and error bars represent the standard deviation. ESM models sizes are represented by colored lines: ESM1v 650M (Orange) and ESM-2 8M, 35M, 150M, 650M, 3B, and 15B (Blues)

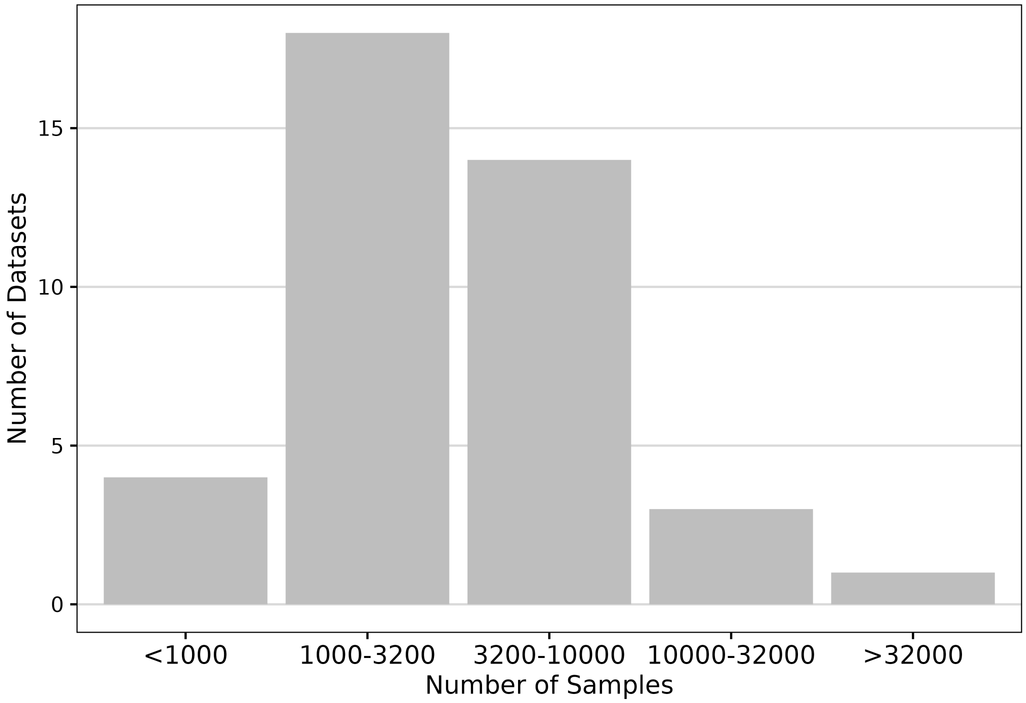


Figure S8: **Sample size distribution in deep mutational scanning (DMS) datasets.**

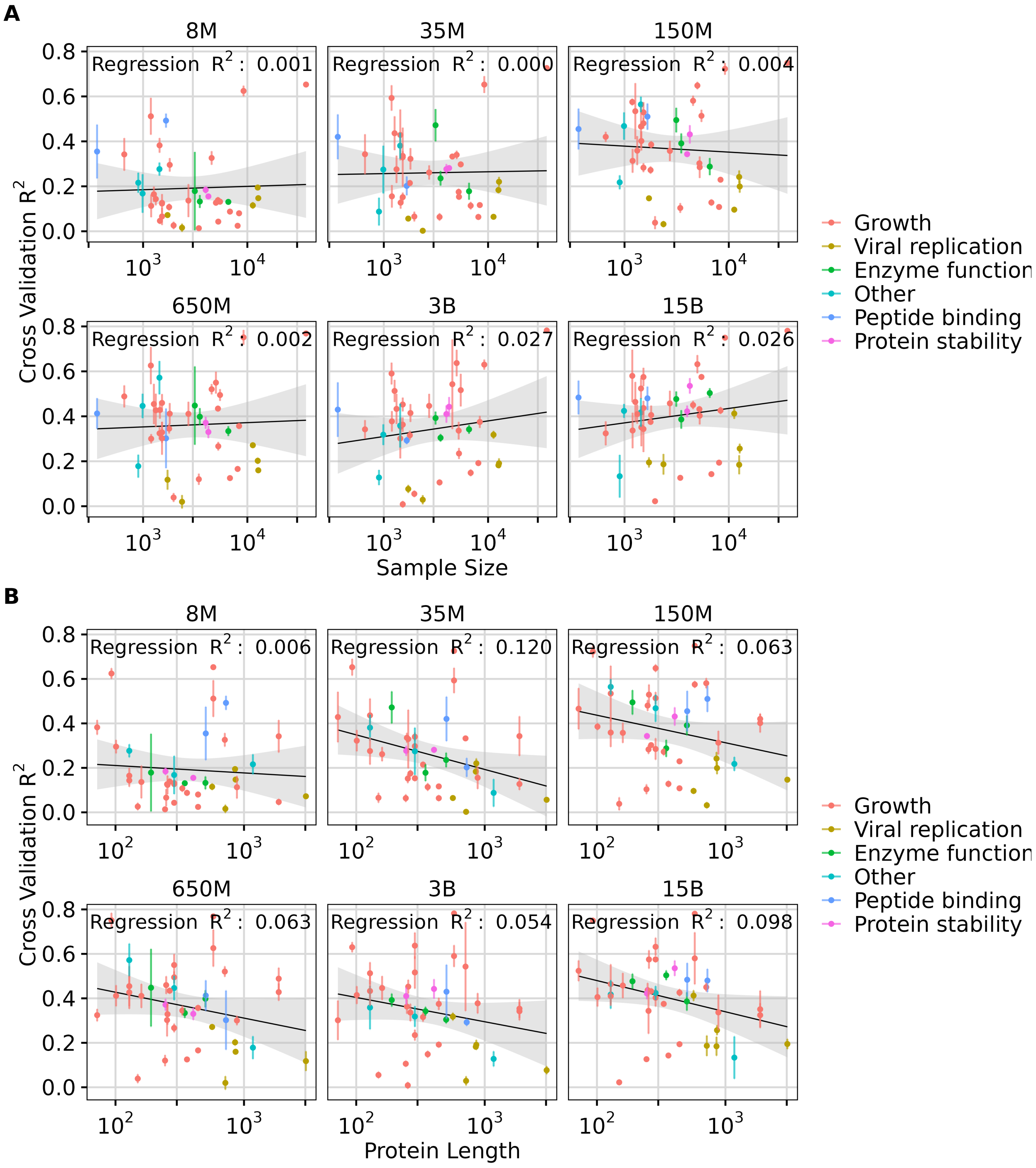


Figure S9: **Effect of sample size and protein length in transfer learning using ESM-2 embeddings.** A) Effect of **sample size** on transfer learning using ESM-2 model embeddings quantified by the regression results ($R^2$ score). B) Effect of **protein length** on transfer learning using ESM-2 model embeddings quantified by the regression results ($R^2$ score). Each dot represents a dataset, and its color indicates the corresponding data measurement type. In each subplot, the y-axis represents the averaged 5-fold cross-validation $R^2$ scores, and error bars represent the standard deviation.

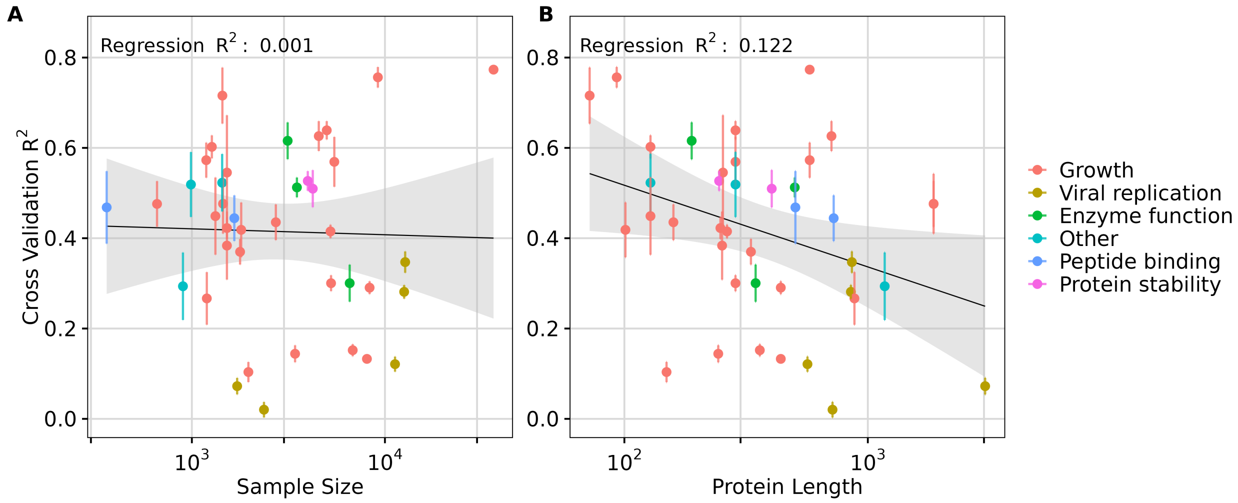


Figure S10: **Effect of sample size and protein length in transfer learning using ESM C 300M embeddings.** A) Effect of **sample size** on transfer learning using ESM C 300M model embeddings quantified by the regression results ($R^2$ score). B) Effect of **protein length** on transfer learning using ESM C 300M model embeddings quantified by the regression results ($R^2$ score). Each dot represents a dataset, and its color indicates the corresponding data measurement type. In each subplot, the y-axis represents the averaged 5-fold cross-validation $R^2$ scores, and error bars represent the standard deviation.

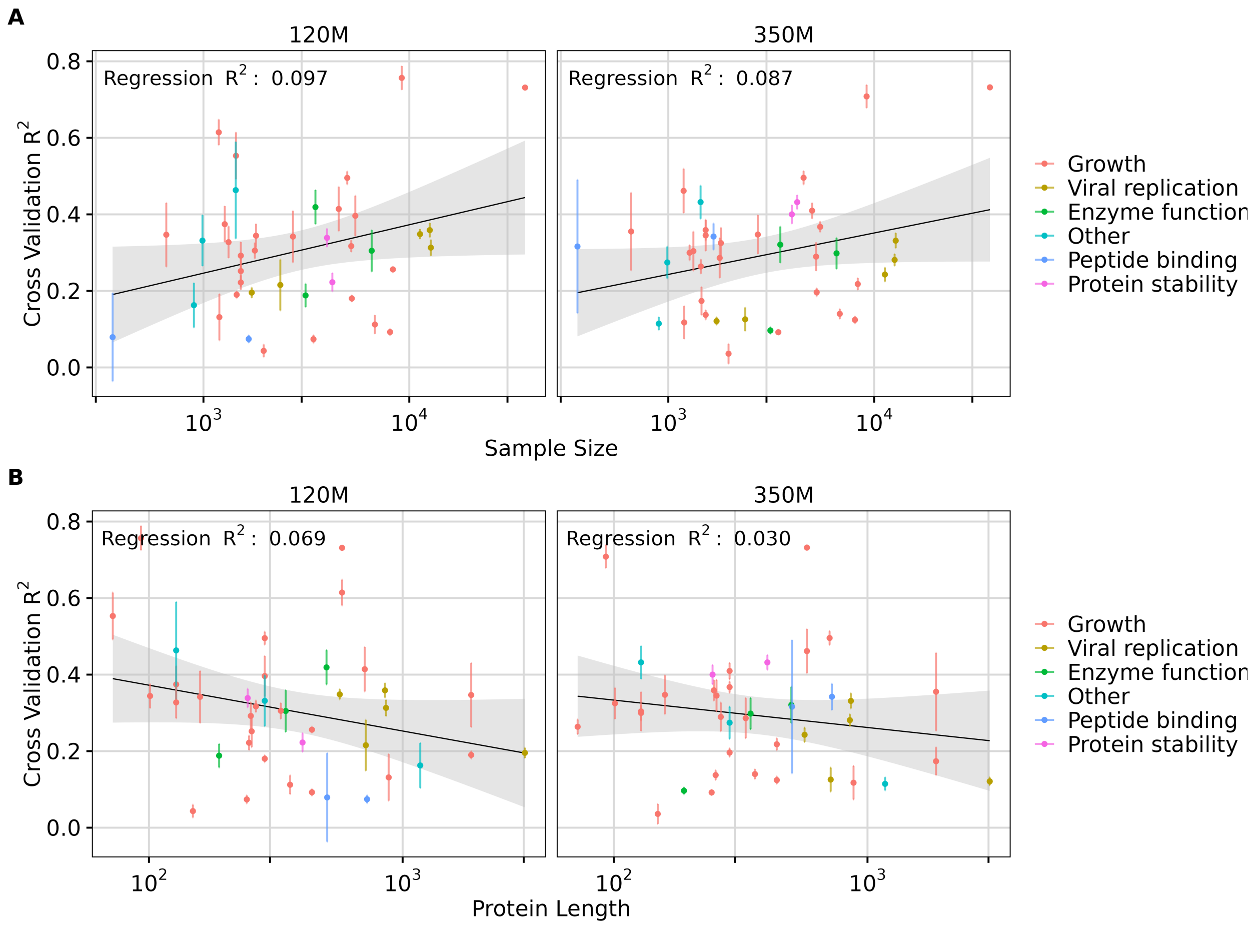


Figure S11: **Effect of sample size and protein length in transfer learning using AMPLIFY embeddings.** A) Effect of **sample size** on transfer learning using AMPLIFY models embeddings quantified by the regression results ($R^2$ score). B) Effect of **protein length** on transfer learning using AMPLIFY models embeddings quantified by the regression results ($R^2$ score). Each dot represents a dataset, and its color indicates the corresponding data measurement type. In each subplot, the y-axis represents the averaged 5-fold cross-validation $R^2$ scores, and error bars represent the standard deviation. The left and right panels represent the two model sizes, 120 million and 350 million parameters, respectively.

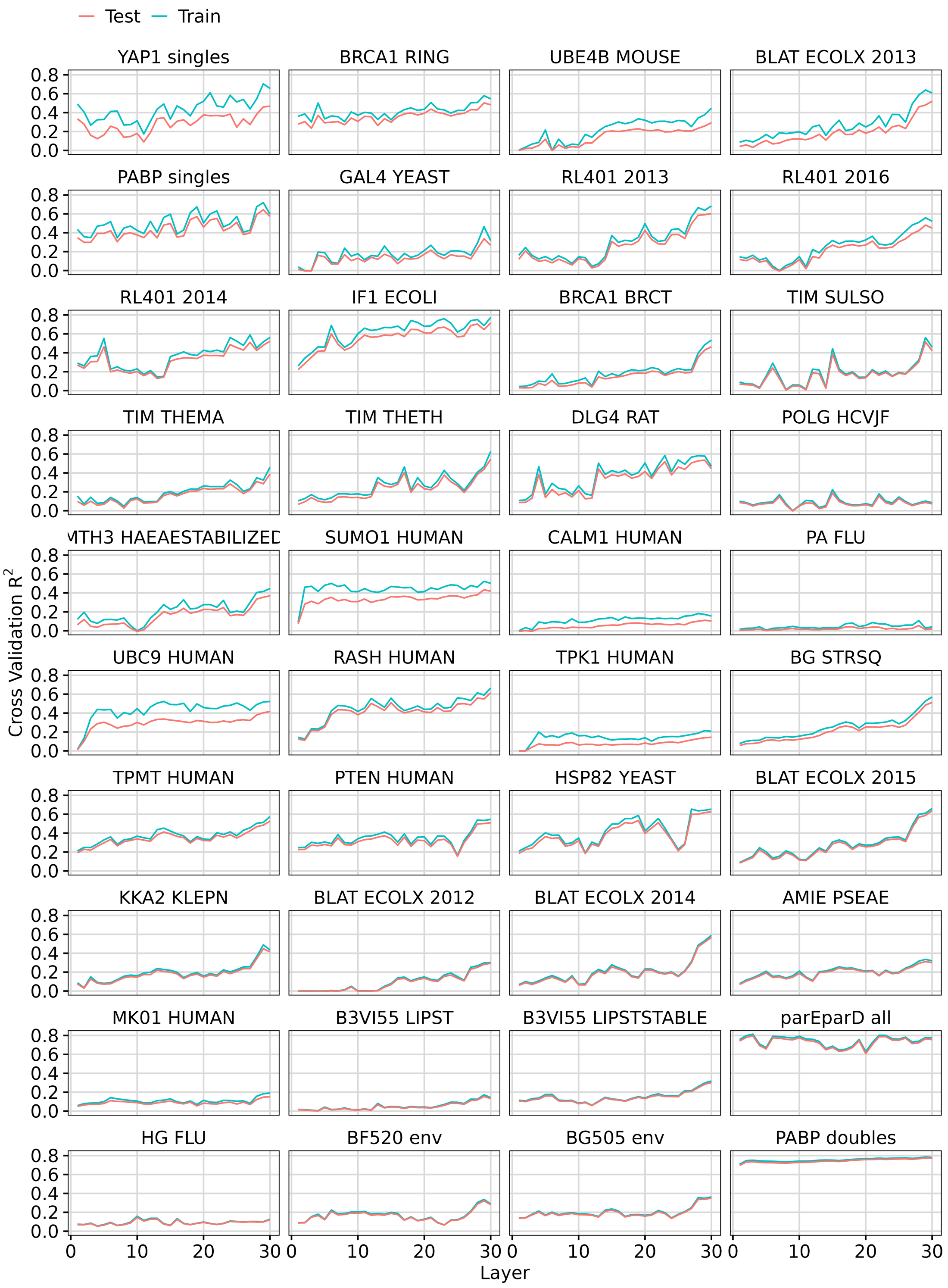


Figure S12: **Performance of Lasso regression using features extracted from different layers of ESMC 300M.** The y-axis represents the average $R^2$ score from 5-fold cross-validation, and the x-axis indicates the layer number.

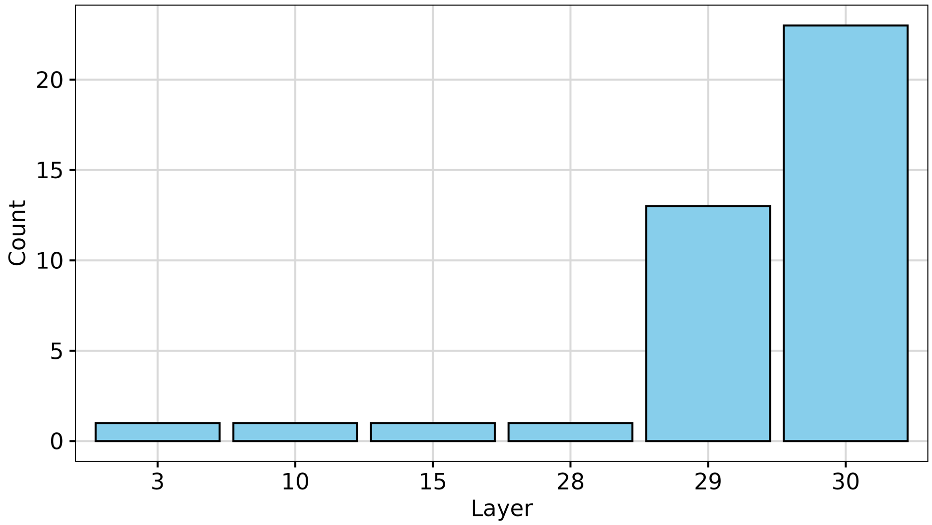


Figure S13: **Cumulative count of datasets for which each layer produced the best performance.** The x-axis represents the layer number used for embedding extraction, and the y-axis shows how many datasets had that layer as the top performer.

Table S1: Target description for PISCES dataset.

| **Class** | **Target** | **Definition** |
|---|---|---|
| Amino-acid Statistics | Alanine | The fraction of alanine residues present in the protein sequence. |
| Amino-acid Statistics | Cysteine | The fraction of cysteine residues present in the protein sequence. |
| Amino-acid Statistics | Leucine | The fraction of leucine residues present in the protein sequence. |
| Physico-chemical Properties (PCP) | Length | Number of residues in the protein. |
| Physico-chemical Properties (PCP) | Charge | The theoretical net charge of a protein sequence, based on the Henderson-Hasselbach equation. |
| Physico-chemical Properties (PCP) | Hydrophobicity | The averaged hydrophobicity values of each residue. |
| Physico-chemical Properties (PCP) | Instability index | The stability of a protein based on its dipeptide composition. |
| Physico-chemical Properties (PCP) | Molecular Weight (Kilodaltons) | The sum of the masses of each amino acid. |
| Physico-chemical Properties (PCP) | Isoelectric point | The isoelectric point (pI), which is the pH at which a particular molecule or surface carries no net electrical charge. |
| Secondary Structure Frequency (SS) | Extended strand (beta strand) | The fraction residues in extended strand conformation. |
| Secondary Structure Frequency (SS) | Alpha helix | The fraction of residues in alpha helix conformation. |
| Secondary Structure Frequency (SS) | Coil | The fraction of residues in coil conformation. |
